# Grid cells show field-to-field variability and this explains the aperiodic response of inhibitory interneurons

**DOI:** 10.1101/101899

**Authors:** Benjamin Dunn, Daniel Wennberg, Ziwei Huang, Yasser Roudi

**Affiliations:** Kavli Institute for Systems Neuroscience / Centre for Neural Computation, NTNU, 7030 Trondheim, Norway; Department of Applied Physics, Stanford University, Stanford, California 94305, USA; Werner Reichardt Centre for Integrative Neuroscience, University of Tubingen, D-72076 Tubingen, Germany; Simons Center for Systems Biology, Institute for Advanced Study, Princeton, New Jersey 08540, USA

**Author notes:** These authors contributed equally to this work.

## Abstract

Research on network mechanisms and coding properties of grid cells assume that the firing rate of a grid cell in each of its fields is the same. Furthermore, proposed network models predict spatial regularities in the firing of inhibitory interneurons that are inconsistent with experimental data. In this paper, by analyzing the response of grid cells recorded from rats during free navigation, we first show that there are strong variations in the mean firing rate of the fields of individual grid cells and thus show that the data is inconsistent with the theoretical models that predict similar peak magnitudes. We then build a two population excitatory-inhibitory network model in which sparse spatially selective input to the excitatory cells, presumed to arise from e.g. salient external stimuli, hippocampus or a combination of both, leads to the variability in the firing field amplitudes of grid cells. We show that, when combined with appropriate connectivity between the excitatory and inhibitory neurons, the variability in the firing field amplitudes of grid cells results in inhibitory neurons that do not exhibit regular spatial firing, consistent with experimental data. Finally, we show that even if the spatial positions of the fields are maintained, variations in the firing rates of the fields of grid cells are enough to cause remapping of hippocampal cells.

## 1 Introduction

Grid cells in the medial entorhinal cortex (MEC) of mammals are selectively active in positions that together form a hexagonal pattern tessellating the space in which an animal moves [1]. These cells, along with other cells in the MEC that code for various spatial quantities, e.g. speed [2], head direction [3, 4], borders [5], as well as hippocampal place cells [6, 7], are believed to form the neural basis of spatial navigation in mammals. Over the past decade, a great deal of knowledge has been collected about the properties of grid cells, thereby gaining better insight into the neural mechanisms involved in spatial navigation [8].

Since the discovery of grid cells, a significant amount of work has been devoted to building models for explaining the underlying neural basis of the spatial selectivity of grid cells [9‒21]. Among these studies, network models based on the concept of continuous attractor dynamics explain the hexagonal firing of grid cells by proposing the generation of a hexagonal pattern of activity on a two dimensional neural array, which is then moved with the animal using velocity input. These continuous attractor models as well as studies on the coding of position by grid cells [22‒26] assume or predict that the different firing fields of a grid cell are periodic replicas of one another, and that the mean firing rate, or amplitude, of the firing fields are similar.

In this paper, by analyzing 373 experimentally recorded grid cells, we first demonstrate that, contrary to the predictions and assumptions in previous work, there is significant field-to-field variability in the response of grid cells, indicating that individual fields of a cell enjoy a significant degree of independence even though they are laid out on a periodic lattice. We then show that the variability in the fields is instrumental in answering one of the problems of existing implementations of continuous attractor models of grid cells: that contrary to experimental findings, they predict regular firing for inhibitory interneurons [27]. Finally, by considering a feedforward network in which summation of grid cell activity of various scales leads to spatially localized activity akin to that of hippocampus place cells, we show that changing the relative amplitude of the fields of grid cells, while maintaining the spatial relationship between the firing of the cells, leads to global remapping, as has been observed in a recent experimental study by Kanter et al. [28].

Continuous attractor models mainly consider a single population of grid cells, and the role of inhibitory neurons is included in the effective interaction between the neurons in this single population [9‒12, 29]. Those cases which do consider separate excitatory and inhibitory populations [13‒15] rely on connectivity rules which result in inhibitory interneurons with grid- or “antigrid”-like spatial tuning. This was shown to not hold true in experimental recordings from the interneurons in the MEC, which did show spatial selectivity but little or no spatial periodicity in the firing patterns of the interneurons [27]. Our analysis of a network of inhibitory and excitatory neurons shows that field-to-field variability together with sparse connectivity between inhibitory and excitatory neurons can explain this experimental finding. Grid cells in the network simulations that we report here express a hexagonal firing pattern through the interacting effect of recurrent connectivity and sparse spatially selective input: some of the neurons are activated by place-like input when the animal visits a certain position, and these neurons activate other cells that should be active at the position via effective inhibitory interactions. The model is thus different from existing continuous attractor models in that the generation of grid activity does not require moving a hexagonal pattern of activity on the network by integrating velocity [9].

## 2 Results

### 2.1 Recorded grid cells show field-to-field variability

We analyzed grid cells recorded from rats with multisite (hyperdrive) implants, reported in Stensola et al. [30]. The cells were recorded from the MEC while the animals explored 1 or 1:5 m square, open-field environments. Analysis was performed per grid cell module. As in [30], modules were identified according to the firing field spacing, orientation and deformation relative to a perfect hexagonal pattern. In total, we analyzed recordings from 373 cells distributed among 13 modules and five rats (see sections 5.1-5.12 for details).

Firing fields were detected in grid cell rate maps via watershed segmentation (see section 5.13 for details). We defined the amplitude of a firing field as the ratio of the number of spikes recorded to the time spent within a circular region enclosing the field.

Although grid cell rate maps often show clear field-to-field variations, as can be seen in figure 1 **a**, it is not obvious whether this variability is mainly induced by some combination of temporal fluctuations in the cell response, stochastic spiking dynamics, and the non-uniform occupancy typically obtained in open-field experiments, or if it is a stable property of the spatial tuning of the cell. We performed a careful statistical analysis to assess the degree of inherent variability in the firing rate of the fields, by measuring the mean firing rate within each firing field, both over the whole experiment and in the first and second halves of the experiment separately. These data are compared, on a cell-by-cell basis, against a synthetic population derived from a spatial tuning profile with identical firing fields, but having the same spacing and spatial phase as the corresponding real cell. Synthetic spike trains were generated from this tuning profile using the real trajectory and assuming Poisson spiking with spatially modulated rate (full details of the procedure is given in section 5.14.) An example of how a real cell and synthetic spikes compare is shown in figure 1 **b**. The figure demonstrates that, when compared to the synthetic spikes, the real cell shows considerably more variation in firing rate from one field to another, and these variations can be substantially larger than the variation from the first to the second half of the experiment. This indicates that field-to-field variations are stable and cannot be explained by occupancy and fluctuations alone.

To make this observation quantitative, we also computed the coefficient of variation c of the amplitudes of each cell, and 1000 synthetically generated spike trains per real cell (see Eq. 25 in section 5.15). Figure 2 **a** shows the measured coefficient of variation of all cells in one module, scattered against the corresponding mean 〈c〉_sim_ computed from synthetic spikes, and compared directly to a single synthetic spike train realization per real cell. The figure clearly shows that the amplitudes of real cells vary much more than those of the synthetic spike trains.

**Fig 1.**
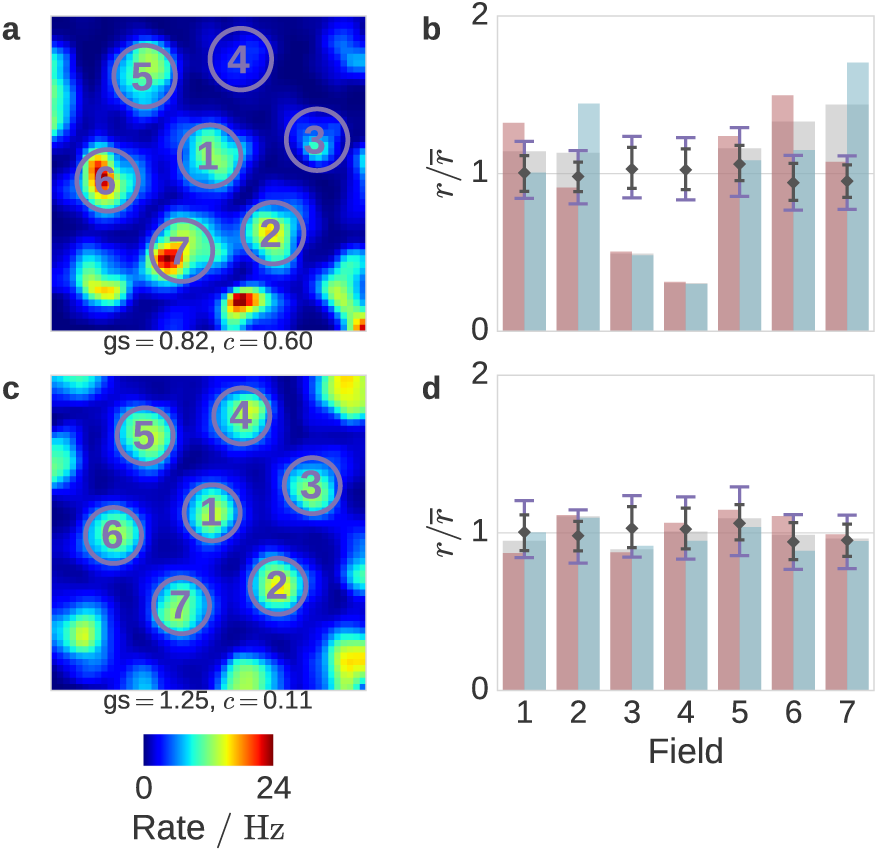
Example grid cell rate map and firing field amplitudes, with simulated counterpart.

**a:** A typical grid cell rate map with detected firing fields circled and numbered. The grid score (gs) and field amplitude coefficient of variation (c) are quoted below the rate map. **b:** The amplitude of each field in **a**, normalized to the mean amplitude of all fields from the same rate map, is shown (gray wide bars in the background) together with the amplitudes measured in only the first and second halves of the session (red and light blue bars in the foreground). On top of this, markers and error bars visualize the distribution of normalized amplitudes from 1000 synthetic spike trains, with the black diamond showing the mean amplitude, the small black whiskers showing the central 95 % of the full-length amplitudes (to be compared with the gray wide bars), and the large purple whiskers showing the central 95 % of the half-length amplitudes (to be compared with the red and light blue narrow bars). For each cell and synthetic spike train, amplitudes are normalized to the mean of the full-length amplitudes from that cell or spike train when creating this plot. **c-d:** Equivalent to **a-b**, but using synthetic spikes generated from the real animal trajectory and a tuning profile consisting of identical firing fields. Another 1000 synthetic spike trains like this were used to generate the error bars in **b** and **c**, and to perform the statistical analysis of the variability.

**Fig 2.**
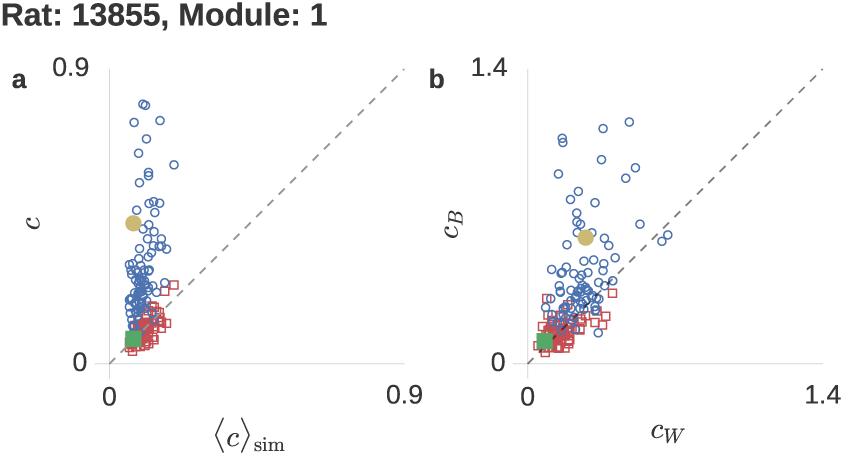
Firing field amplitude variability.

**a:** Blue circles show the field amplitude coefficient of variation of each cell in a module, scattered against the mean coefficient of variation from the 1000 synthetic spike trains corresponding to that cell. Red squares show the same quantity for one particular synthetic spike train per real cell. **b:** Between-field coefficient of variability scattered against within-field coefficient of variability from the same real cells and synthetic spikes as in **a**. Variabilities are computed from field amplitudes measured in the first and second halves of the experimental sessions separately. For each cell, we only included the amplitudes of firing fields that were detected in both the real cell rate map and all corresponding synthetic rate maps. If less than 3 such fields were found, the cell was discarded. The enlarged, filled markers in **a** and **b** correspond to the cell (yellow circle) and synthetic spikes (green square) shown in figure 1.

While this comparison gives a first indication of the degree of variation in grid cell field amplitudes, it does not take into account the possibility that the measured field amplitudes in a finite-time experiment may be affected by factors not accounted for by the synthetic spike trains, such as the simultaneous encoding of other kinds of information in the grid cell (i.e. head direction), as well as non-Poisson and possibly temporally varying spiking statistics. We therefore considered data from the first and second halves of the experiments separately, and asked whether amplitude variation between fields exceeded what would be expected based on the variability over time of the amplitudes of the same cell. Figure 2 **b** shows the between-field coefficient of variability *c_B_* scattered against the within-field coefficient *c_W_* for each cell in a module, as well as for a single synthetic spike train per real cell. The coefficients *c_B_* and *c_W_* are computed from between-field and within-field variabilities, quantities that are standard in one-way ANOVA (see section 5.15 for details). Hence, a *c_B_* that is significantly greater than *c_W_* indicates that we see more variation between the field amplitudes than what would be expected based on the temporal fluctuations we observe. From the figure we see that this is the case for a large fraction of the real cells, but certainly not all. On the other hand, all the simulated cells have *c_B_* and *c_W_* of comparable magnitudes and somewhat smaller than the typical values for the real cells. Figure 1 **b** shows what this can look like in terms of actual field amplitudes: the difference between the first and second half is quite large for certain firing fields when compared to the synthetic data, but still much smaller than the variation between different fields in the real data.

To condense this notion into a single number, we computed the F-statistic 
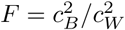
. We used the 1000 synthetic spike trains to generate a cell-specific null distribution for the F-statistic, which was then used to compute a p-value by ranking the real F-statistic against this distribution. Note that although we once again invoke the synthetic data here, we only compare between-field variabilities to within-field variabilities from the same spike train, such that the effects of factors not accounted for in the synthetic spike generation should cancel out to a first approximation. Thus we expect that, regardless of additional factors affecting the between-field variation, each generated distribution for the F-statistic will predict with reasonable accuracy the outcome for a real cell with this particular trajectory, firing field locations, and mean firing rate, assuming that the firing fields are identical in terms of the strength of the underlying spatial tuning. This assumption could be rejected with *p* < 0:05 for 24 of the 86 cells in the module shown in figure 2, and 129 of the 373 cells in total.

The fact that only a little more than a third of the cells have a statistically significant between-field to within-field variability ratio reflects a continuum in the cell population, from cells where this ratio is of order unity to cells to cells with very large field amplitude variations. This property is also clearly visible in figure 2. However, the number of statistically significant cases greatly exceeds the chance level (i.e. 5 % of the population). By interpreting rejection or not of the null hypothesis as a Bernoulli trial with success probability 0:05, the number of rejections in a population of cells should follow a binomial distribution under the null-hypothesis, and thus we can define an aggregate *p*-value *p*_tot_ for the number of rejections in the population. For the module shown in figure 1, this comes out to *p*_tot_ = 3.5 · 10^−12^, and for all cells in the data set we find *p*_tot_ = 7.5 · 10^−71^. This leads us to conclude that the discrete translational symmetry of the grid cell network with respect to physical space is broken not only by geometrical distortions of the lattice due to the environment [31], but also in terms of the firing field strength at each site.

Supplementary figure 4 contains figures similar to figure 2 for all modules in the data set, containing between 2 and 93 cells after discarding cells for which fewer than three firing fields were detected. On average, about 35 % of the cells in a module show statistically significant amplitude variations. We find *p*_tot_ < 0.05 for 9 of the 13 modules, and *p*_tot_ < 0.005 for 8 of these.

### 2.2 Basic aspects of the recurrent excitatory-inhibitory models

Networks models of grid cells based on continuous attractor dynamics and integration of velocity involve two key steps. The first one is to generate a translationally periodic pattern of activity in a network of neurons, and the second is to translate this pattern along with the animal movement using speed and head direction input [9]. The fact that the fields of a single grid cell appear due to the translation of a regular pattern, guarantees that in these models each firing field of any given grid cell is the same as another field of the same cell. The analysis in the previous section thus demonstrates that this simple mechanism cannot be the whole story and other mechanisms are in play to generate the field-to-field variations in the firing rates of a grid cell.

As described in the Introduction, recent experimental evidence also shows that inhibitory neurons in the MEC do not exhibit spatially periodic firing [27], as predicted by existing continuous attractor models. In this section we consider a two population network of excitatory and inhibitory neurons and show that, with appropriate connectivity, field-to-field variability can lead to irregular firing, and that this comes at the cost of the ability to integrate velocity properly.

We consider a simple two-population models of the form [32]:

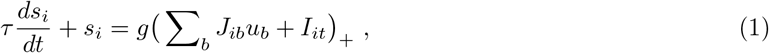

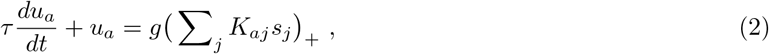

where *s_i_* and *u_a_* represent the firing rates of the excitatory neuron (grid cell) *i* and inhibitory neuron (interneuron) *a*, respectively. (…)_+_ is the threshold-linear function, which outputs zero if its argument if equal to or less than zero and is a linear function for positive arguments. *g* is the single neuron gain, *τ* the neuronal time constant, *J_ib_* and *K_aj_* the connectivity from interneurons to grid cells and grid cells to interneurons, and *I_it_* the external input.

We consider different forms of external input *I_it_* during the simulations reported below. Classically in continuous attractor networks of grid cells, one considers a velocity dependent of the typical form

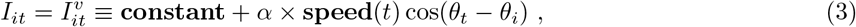

where the superscript *v* emphasizes the velocity dependence of the input, speed(*t*) and *θ_t_* are the time-varying speed and direction of the animal as it moves around, the velocity modulation and *θ_i_* the preferred direction of neuron i, patterned as in Burak and Fiete [11] to (0, π/4, π/2, 3π/4).

However, as noted in the Introduction, with pure velocity input we cannot see any variability in the firing field amplitudes of grid cells: even if the connectivity is heterogeneous such that in the stationary pattern the peak activity at some positions in the network ends up being different from another one, the translation of the pattern will lead the fields of each grid cell to have exactly the same amplitudes, with the heterogeneity only causing the rates being different from one cell to another, and not different within one cell. We therefore consider the case in which the hexagonal firing pattern and the variability in the fields of one cell arise from a different type of input to the excitatory neurons in the model, the place-like input, that could originate from a combination of sensory input as well as the hippocampus or other areas with spatially selective neurons projecting to the MEC. Assuming a population of place-like neurons whose rates are denoted *P_j_(t)*, we take the input to be of the form

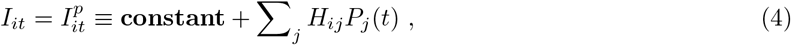

where the superscript *p* indicates the place-like selectivity of the cells contributing to this type of input, and *H_ij_* are weights that connect the place-like neurons to the excitatory grid cell population. The weights are chosen from place-like inputs that are active at hexagonally arranged fields in the environment. The amplitude of these *H_ij_* are drawn from a Gaussian distribution of mean *μ* = 0.5 and standard deviation σ with negative values set to zero, as suggested in [33]. We also dilute the input by setting a fraction, *f_d_*, of these connections to zero, allowing us to study not only the effect of variability in the amplitude of the place-like input but also the sparsity of the input. In what follows, we will study the effect of σ and *f_d_* on the quality of the grid cells generated by the model as well as the field-to-field variability. But before doing that, we discuss in the next section two ways of wiring up the neurons in the model, namely the choices for *J* and *K* in Eq. 2.

Examples of networks with inputs of the form in equation 3 and 4 can be seen in accompanying videos 1 and 2.

### 2.3 Two ways of generating the connectivity matrices

Having described the types of input to the network that we consider, in this section we detail two ways of designing the connectivity pattern between neurons in the model. To do so, we first note that as *du_a_*/*dt* → 0, we have 
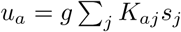
since *K_aj_* is non-negative and g(.)+ is a zero-threshold linear activation function. We can therefore approximate the term 
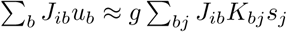
 to end up with the following effective connectivity between the excitatory neurons:

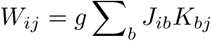

As a simple effective connectivity, similar to that used in the literature [11,12], we let

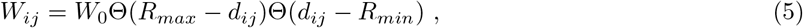

where Θ(·) is the Heaviside step function, *R_max_* the outer ring of the radial extent of the connectivity, *R_min_* the inner ring, *W_0_* the strength of the inhibitory connections and *d_ij_* the Euclidean distance between cells *i* and *j*.

For the simulations reported below, where the external input includes the velocity terms the Euclidean distance, *d_ij_* is biased in the preferred direction *θ_i_* of the grid cell [11],

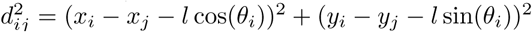

with *x_i_* ∊ [1,*N_x_*], *y_i_* ∊ [1,*N_y_*] representing the position of neuron *i* in a two dimensional *N_x_* × *N_y_* neural sheet with periodic boundary conditions and spatial offset *l*. Each grid cell in the model is therefore connected with other neurons within a radius between *R_min_* and *R_max_*, forming an inhibitory ring connectivity. For simulations where place-like input is used, no bias exists in the connectivity.

Our two network architectures now amount to the different ways of choosing the connectivity matrices **J** and **K** to obtain the prescribed effective connectivity **W**. In principle, this problem is under-constrained and has an infinite number of solutions. Conveniently, it has been shown that grid cells in superficial layers of MEC are primarily connected through inhibition [12,13], making the matrix *W_ij_* strictly non-positive while also justifying the lack of direct connectivity between the excitatory neurons *s_i_*. Furthermore, by Dale’s law, is strictly non-negative and *J_aj_* is non-positive, hence we rewrite the factorization problem as (−*W*) = *g*(−**J**)**K**, where all three matrices are non-negative.

We considered two possible ways of generating the connectivity. First, we initialized both **−J** and **K** to random numbers drawn from a uniform distribution on [0; 0.1] and solved for these iteratively using methods for factorizing non-negative matrices (NMF) [34,35]. NMF is an iterative method that converges to a local minimum of the cost function 
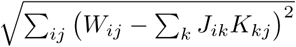
 (see section 5.16-5.17 for additional details). This minimum is not necessarily the global minimum, so different initializations may result in different connectivity patterns. We refer to this approach as the fully factorized model. For the second approach, which we label the partially factorized model, we set the connections to the interneurons **K** to sparse (10 %) random excitatory connectivity, and solved for **J** only using NMF.

Figure 3 shows examples of the connectivities that we find with these two approaches. As can be seen in figure 3 **a,b,c,** both approaches manage to reasonably approximate the desired effective connectivity. In the case of the fully factorized model, this resulted in interneurons that receive inputs from grid cells of similar spatial phase (figure 3 **d**) or receive input from grid cells of surrounding spatial phases (figure 3 **e**). These interneurons will exhibit grid-like or anti-grid like firing correspondingly, similar to those in [13] and [14]. These grid-like and anti-grid-like interneurons then project back to the surrounding (figure 3 h) and center (figure 3 **i**), respectively. The resulting connectivity for the partially factorized model (figure 3 **c**) also manages to approximate the desired connectivity, but typically requires many more interneurons than the fully factorized model, e.g. in the example shown it required 5 times more interneurons than excitatory neurons. This can be attributed to the fact that in the fully factorized model both inhibitory-to-excitatory connections and excitatory-to-inhibitory connections can be adjusted to achieve the desired connectivity while in the partially factorized one only the former are allowed to be adjusted. In the partially factorized model, the inhibitory neurons obviously receive random input from the grid cells (figure 3 **f,j**) while projecting back in a non-random but not obviously structured manner (figure 3 **g,k**). Consistent with this connectivity scheme, Buetfering et al., 2014, found that grid cells of all phases functionally project to local interneurons [27].

Using these two approaches we were able to construct different models with variable numbers of interneurons while maintaining the same effective weight matrix *W_ij_* between the grid cells. Additional details on the construction of the grid cell model can be found in 5.16.

**Fig 3.**
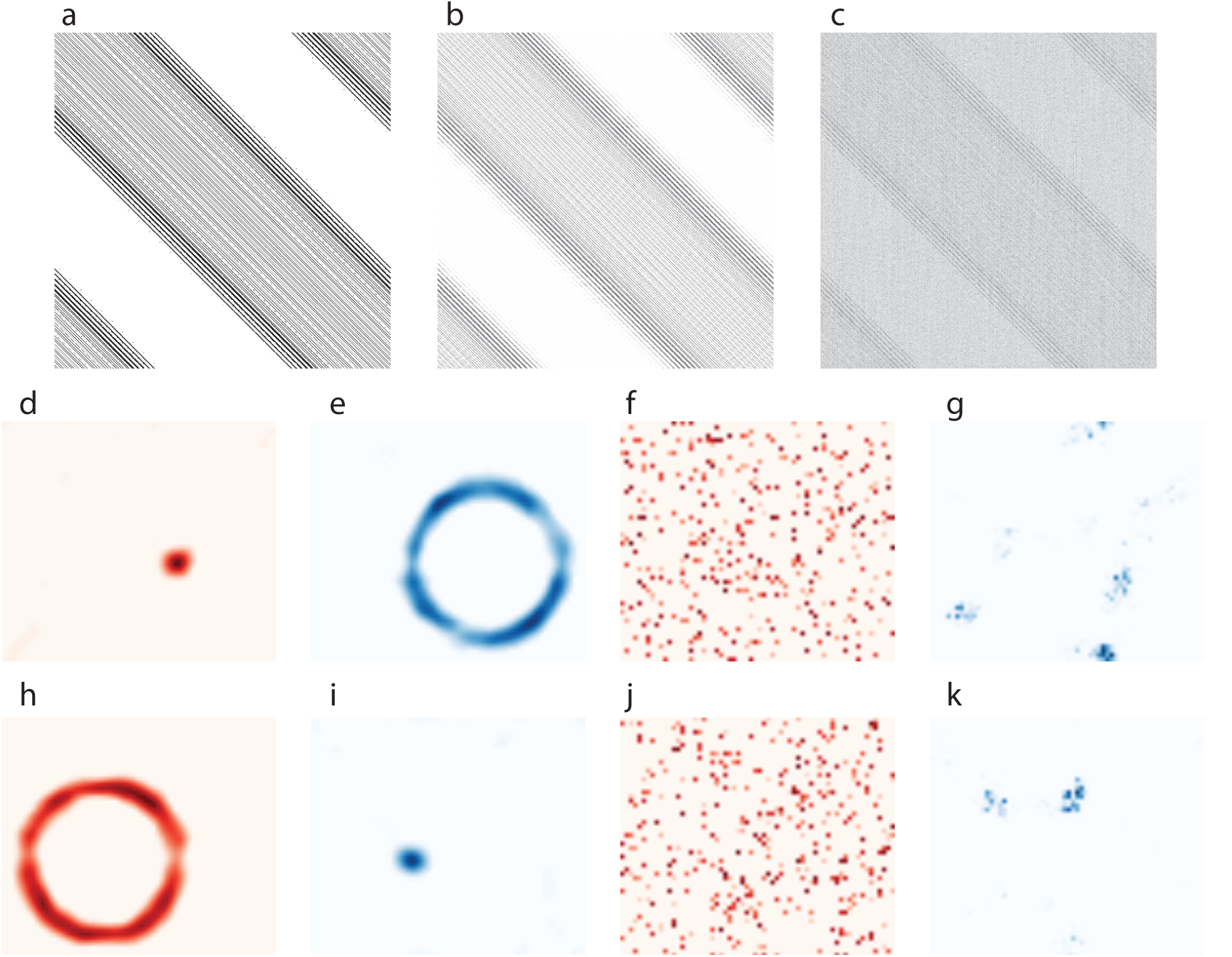
Approximate connectivity using non-negative matrix factorization.

(**a**) The desired effective connectivity matrix between grid cells, *W_ij_* in Eq. 5, with σ, *f_d_* and offset, *l*, equal to zero. (**b**) the resulting effective connectivity for the fully factorized model, with 200 interneurons and 896 excitatory cells and (**c**) for the partially factorized model, with 896 excitatory and 4480 inhibitory cells. (**d,h**) Examples of the connections from the 32 × 28 excitatory cells arranged on a two dimensional lattice to two example interneurons in the fully factorized model and (**e,i**) the corresponding inhibitory-to-excitatory connectivity from the same interneurons to the excitatory neurons. In each of the examples the aptitude of the connections are normalized to the peak value of the coupling matrix with excitatory couplings colored red and inhibitory blue. In **d,e,h,i** the excitatory cells are arranged according to spatial phase. The interneuron in **d** receives input from excitatory cells with similar phase but and inhibits a number of excitatory cells e. Since this inhibitory neuron is activated by grid cells with similar phase, it is going to exhibit gird-like firing. The opposite is true for the example inhibitory neuron **h,i** which will show anti-grid-like tuning. **f,g,j,k** Same as **d,e,h,i** but for the partially factorized model.

### 2.4 Response of models to velocity modulated and place-like inputs

We subjected both models to input of the form shown in Eq. 4, following an experimentally recorded trajectory of an animal exploring a square open-field environment. In both models, we could change the degree of firing field amplitude variance in the excitatory grid cells by changing the dilution *f_d_* of the place-like input as well as the standard deviation σ of the truncated Gaussian distribution from which the connections *H_ij_* in Eq. 4 are drawn. The fields of a grid cell that were not directly excited by place-like input added further variations in the field-to-field difference of the cell: a field not driven by place-like input would be activated only through the interaction between the external excitatory drive that all neurons receive (i.e. the constant term in Eq. 4) and the pattern forming effective inhibitory connectivity, leading to a lower firing rate compared to fields driven by place-like input.

In figure 4, we show how the firing properties of the cells change as we vary *σ* with fixed *f_d_* = 0.2 for both fully factorized and partially factorized models, by showing rate maps and cross-correlograms of selected cells.

**Fig 4.**
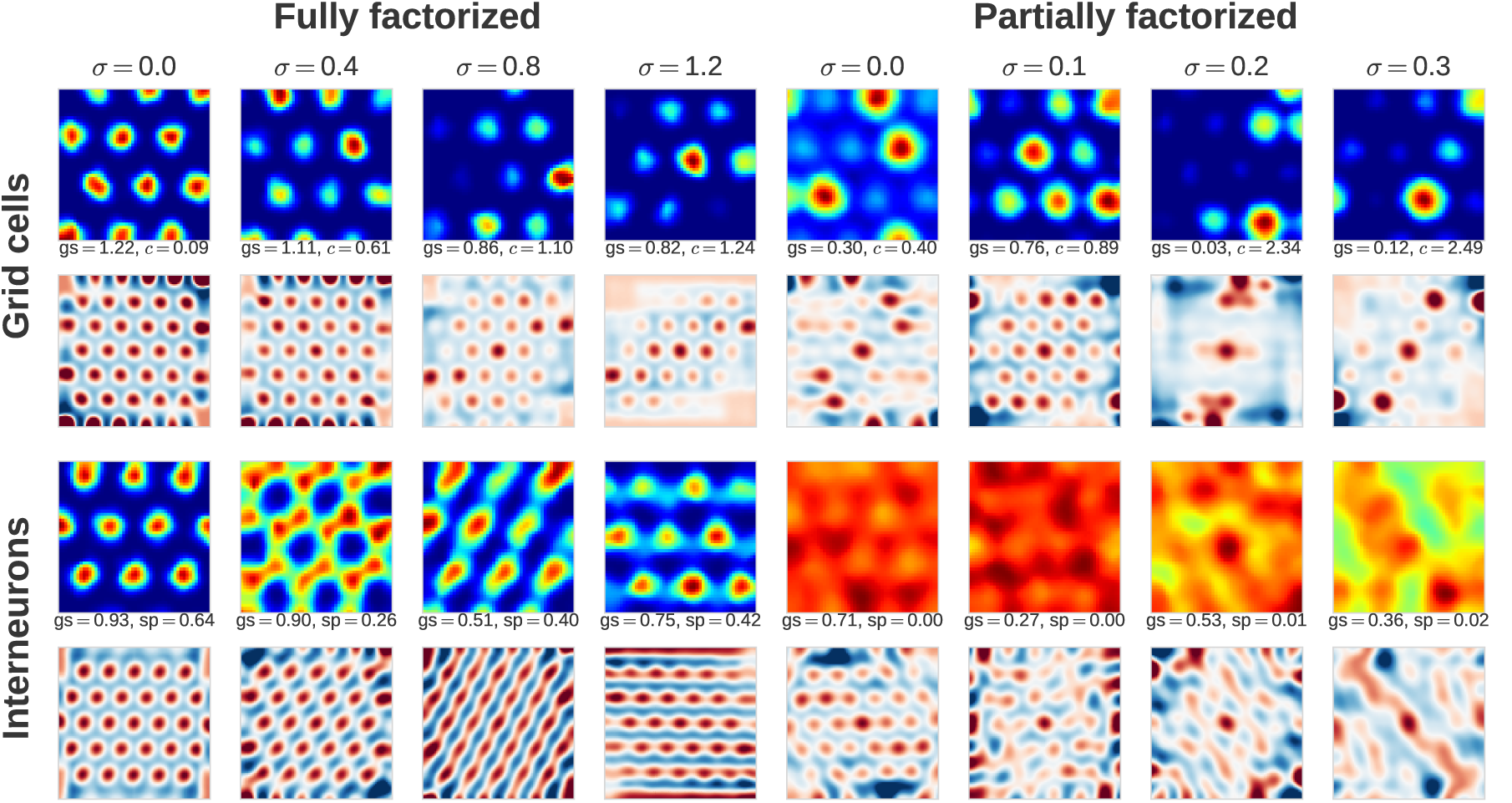
Rate maps and autocorrelograms of simulated neurons. Rate maps and autocorrelograms of one excitatory and one inhibitory neuron from each of several model simulations with different parameters. The top two rows show excitatory neurons (grid cells), with grid score (gs) and field amplitude coefficient of variation (c) quoted between the rate map (above) and autocorrelogram (below). The bottom two rows show inhibitory neurons (interneurons), listing grid score (gs) and sparsity (sp). The four left columns are taken from runs of the fully factorized model with different values of place-like input connection strengths σ, and the four right columns from similar runs of the partially factorized model with dilution *f_d_* = 0.2. Complementing the observations in figure 5, we see that inhibitory neurons in fully factorized models have a high degree of periodicity regardless of the connectivity variance, while interneurons in the partially factorized model lose their periodicity as the grid cell amplitude variability increases. Note that inhibitory neurons in the fully factorized model need not have isolated firing fields even though they are periodic; see Supplementary figure 6 for several counterexamples.

As can be seen, in the fully factorized model, the excitatory neurons but not the inhibitory neurons become more irregular as σ is increased. This is, however, not the case for the partially factorized model. In this model, both inhibitory and excitatory neurons become more irregular as σ is increased.

In figure 5, we show the trends exemplified in figure 4 at the population level. As can be seen in figure 5 **a-b**, the grid score for the excitatory neurons but not the inhibitory neurons drop as σ is increased in the fully factorized model. In fact, there is no value of σ where, in this model, we can see grid cells with grid score similar to the real data and irregular inhibitory neurons (see also figure 4). This is, however, not the case for the partially factorized model. In this model, the grid score for both inhibitory and excitatory neurons drop by increasing σ. In fact, for the parameters we chose already at σ close to zero and dilution level *f_d_* = 0.2, we can have interneurons with irregular firing (figure 4) while having excitatory neurons with realistic grid score. As we show in figure 5 **c**, the irregularity in the fields can also be increased by keeping σ at zero and simply varying the dilution *f_d_* of the place-like input.

The grid score on its own suffers from the fact that it can be low due to field strength variations even if the pattern is regular (see also figure 4). We therefore also analyzed our results using a modified score which is a linear sum of the grid score as well as the spatial sparsity of the firing, which we will denote the *combined score* (see section 5.5 for further details). For grid cells most of the firing is confined to isolated fields, so taking into account the spatial sparsity helps separate such cells from e.g. interneurons, which may exhibit some degree of periodicity but often fire all over space. As can be seen in figure 5 **d-f**, in the fully factorized model both grid cells and interneurons maintain high scores, while the interneurons of the partially factorized model have substantially lower scores. Inspection of the firing rate maps of individual cells (see Supplementary figure 6 for more examples) clearly substantiates the conclusion from this figure that in the

**Fig 5.**
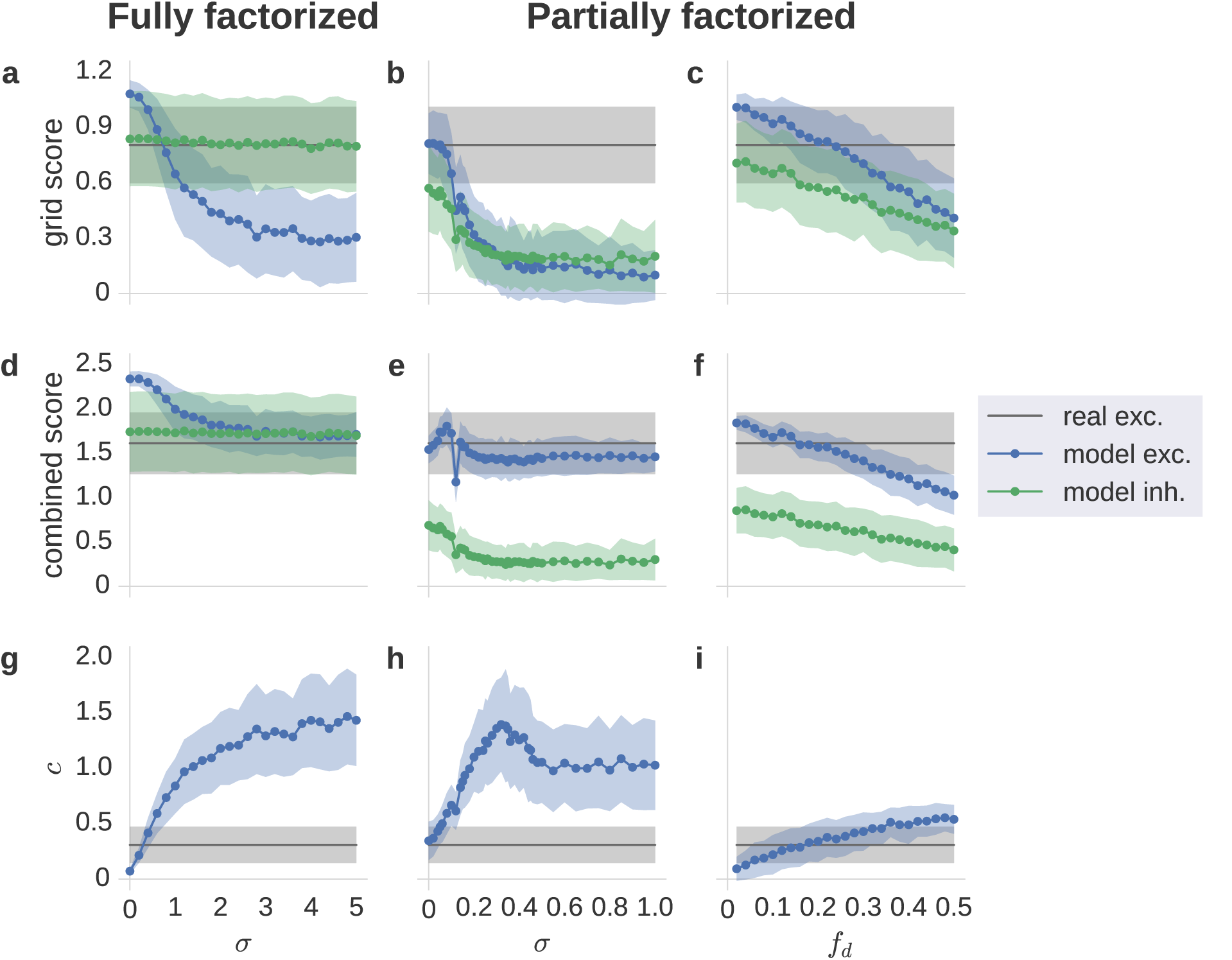
Connectivity variance and the resulting selectivity of excitatory and inhibitory neurons. **a-b:** Mean and standard deviation of the grid score of excitatory grid cells (blue) and inhibitory interneurons (green) from several model simulations, plotted against the standard deviation in the place-like input connection strengths, *σ*. For comparison, the same quantities for real grid cells are shown in gray. **c**: Same as **b** but keeping **σ** fixed and varying the dilution fraction *f_d_* of non-zero couplings randomly removed. **d-f**: Mean and standard deviation of the combined score of grid cells and interneurons from model simulations, and real grid cells. **g-i**: Mean and standard deviation of the firing field amplitude coefficient of variation c of excitatory grid cells from model simulations, and real grid cells. **a,d,g**: Fully factorized model. Grid scores and combined scores of interneurons remain high and similar to real grid cells for all connectivity variances, reflecting their periodic and relatively sparse firing in this model. Grid scores of grid cells decrease with increasing input variance, but only due to increasing field amplitude variation, not geometric distortions of the firing pattern (see figure 4). **b,e,h**: Partially factorized model with *f_d_* = 0.2. Grid scores of model grid cells are similar to real data for zero input variance, and decrease with increasing input variance. In this model, grid scores of interneurons also decrease as σ is increased, reflecting their waning periodicity (see figure 4). Due to their non-sparse firing, interneurons are clearly separated from grid cells by the combined score. **c,f,i**: Partially factorized model with σ = 0. Without dilution (f_d_ = 0), both the interneurons and grid cells are predictably periodic with little field-to-field variations. By diluting the projections from the place-like input, the field amplitudes of the grid cells become increasingly variable, while the grid and combined scores of the interneurons decrease as in **b,e,h**.

fully factorized model, increasing the variance of the input does yield variable field amplitudes in the grid cell but the inhibitory neurons maintain a high degree of regularity and sparsity, while for the partially factorized model, the inhibitory neurons have non-sparse firing, and also lose their regularity when the variance of the input to the excitatory neurons increases. Note that, as shown in figure 5 **g-i**, in both networks the coefficient of variation in the field-to-field firing in the excitatory neurons can be reached at small values of σ and *f_d_*, but as described above, it is the behavior of the inhibitory neurons that sets the two architectures apart from each other.

### 2.5 Integrating velocity

After establishing that with the place-like input both the fully factorized and the partially factorized models generate grid cells, we wondered whether these two architectures can also generate grid cells using only velocity-head directional input (Eq. 3). While the fully factorized model managed to generate hexagonal grid cells only using input of the form of Eq. 3, the partially factorized model failed to do so, e.g. figure 6. Testing various configurations with up to 20 times more interneurons than grid cells we were unable to find a configuration that could accurately integrate the velocity and generate grid cells in the partially factorized model. As can be seen in figure 6 (right), with a ratio of interneurons to grid cells that is completely biologically implausible, the model was only able to produce random, unreliable firing fields when attempting to integrate over velocity for ten minutes. At the same time, the fully-factorized model performed beyond expectations: with as little as 200 interneurons (comprising approximately 5 % of the network) and 64 × 56 grid cells, the model could accurately path integrate over 10 minutes of tracked position data (figure 6, left).

**Fig 6.**
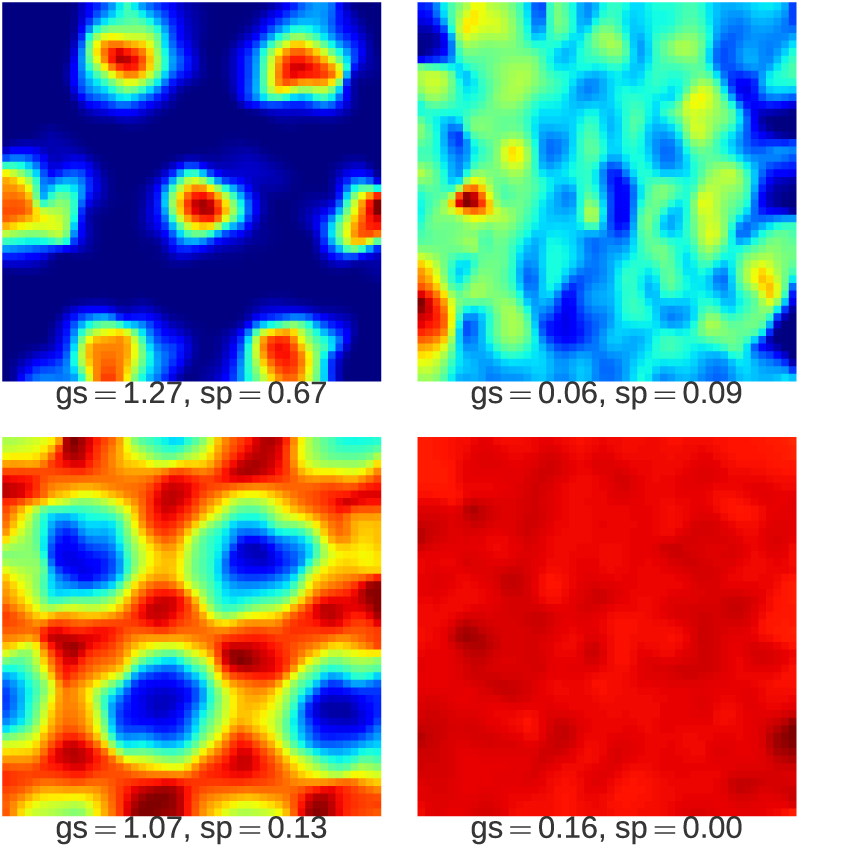
*Top left*: Rate map of a simulated grid cell arising from stable path integration of velocity input in the fully factorized model. *Bottom left*: Rate map of a simulated interneuron during the same simulation run. *Top right*: Rate map of a simulated would-be grid cell that fails to develop a grid pattern because path integration is too unstable. *Bottom Right*: Rate map of an interneuron during the same simulation run. No discernible grid-like structure was observed after a ten minute session.

To understand why the two models performed so differently under path integration, we considered the issue of drift [36]. Since even minute levels of randomness in connectivity is known to cause drift, drift is a likely candidate for the failure of the partially factorized model for generating grid cells by integrating velocity because of the presence of random sparse connectivity in this network. For each drift simulation the pattern was initialized randomly, with the external drive *I_t_* set to a constant value, allowing the network to converge to a static pattern. Once formed, the pattern was shifted randomly around the neural sheet. In an idealized continuous attractor, the shifted pattern should stay in place indefinitely due to the translational invariance property [36], but any inhomogeneity in the networks, makes the pattern prone to drifting. We measured the drift as the average distance traveled from 200 shifted locations during 100 time steps, allowing us to estimate the average drift velocity during the first second after shifting the pattern (see figure 7).

For the fully factorized models, the resulting drift decreased exponentially with the number of interneurons included in the model (see figure 7) while the partially-factorized model decayed significantly more slowly, requiring several orders of magnitude more interneurons to achieve a similar degree of stability.

**Fig 7.**
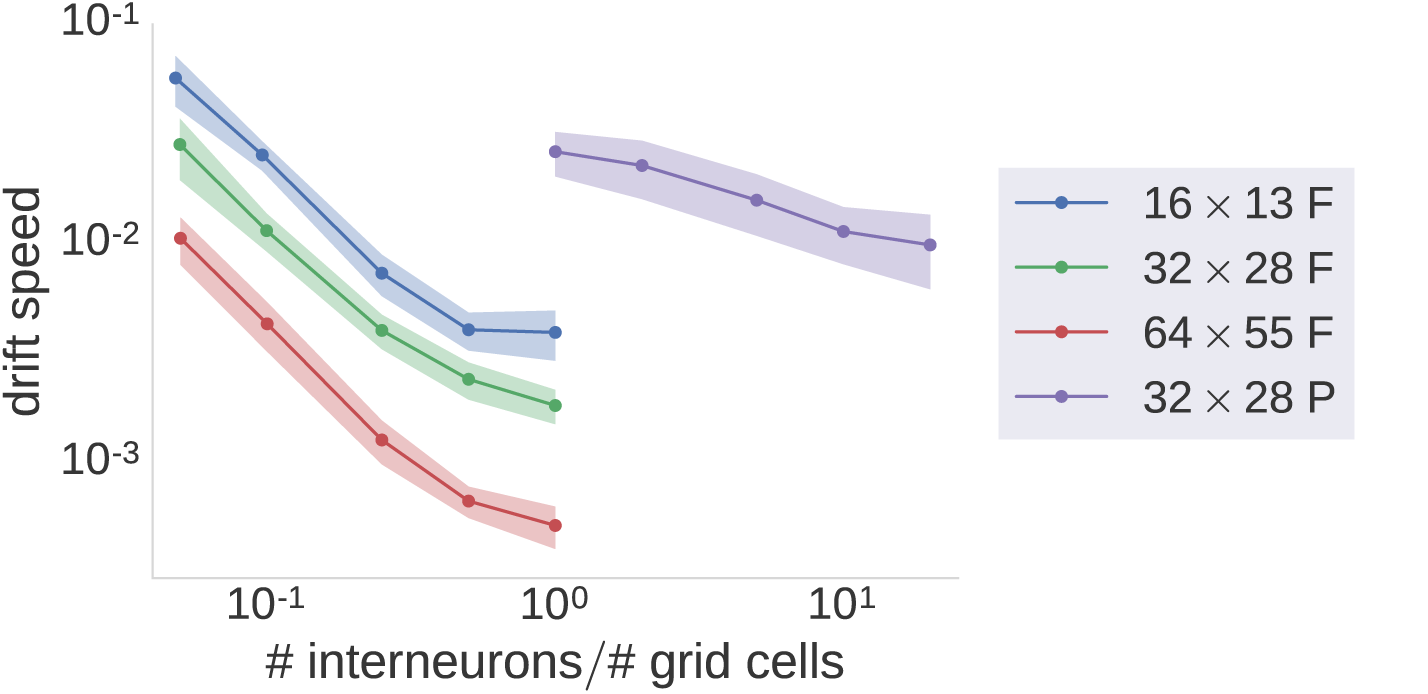
Stable continuous attractor dynamics. Mean and standard deviation of the measured drift speed (*y*-axis), defined as (displacement on neural sheet/(time step · length of neural sheet). The drift speed is plotted against the ratio of the number of interneurons to the number of grid cells (*x*-axis) for various configurations of both the fully (F) and partially (P) factorized model. The legend shows the size *N_x_* × *N_y_* of the neural sheet.

## 3 Field variability and remapping

The fact that the firing rate of individual fields of a cell may shows strong and robust variations may have multiple implications. One is that it will increase the information that the firing of each individual cell carries about the position of the animal: if all fields are identical, it is impossible to distinguish between different sites by observing the firing of the cell, and this ambiguity could be reduced by introducing varying field amplitudes. This could have implications for theories on how grid cell parameters are chosen for optimally encoding space [37,38].

In addition, variability among firing fields suggest the possibility of an additional role for grid cells in distinguishing between different environments, i.e. encoding contextual information. Previous work assuming equal firing amplitudes [39] confirm the high spatial information content of grid cells while suggesting that they provide almost no contextual information (that is, they cannot be used to discriminate between environments). Interestingly, a recent study by Kanter et al. [28] shows that with a chemogenetic manipulation of the mouse MEC, random shifts in the firing amplitudes of the individual grid fields can be induced without changing the positions of fields. This observation can be exploited in a simple feedforward model in which the output of grid cells with different spacings are summed and thresholded by cells in the hippocampus, such that the hippocampal cells obtain spatial tuning [22]. With identical fields, the only way to obtain remapping in the place cells is to change the relative spatial phase of different grid cells. With non-identical fields, however, changing the relative amplitude of different fields can cause remapping to occur, as shown in figure 8. The statistical analysis and theoretical model presented here, together with this recent experimental finding, thus suggests an expanded role for grid cells, not only integrating over velocity output, but over spatial and contextual cues more generally.

## 4 Summary and Discussion

In this paper we established that the firing fields of real grid cells are not simple translations of one another, but that their amplitudes can in fact vary significantly. These variations are stable and reproducible, and are not merely a consequence of experiments with finite duration and imperfect, non-uniform sampling of the environment, or temporal fluctuations in the spiking of the neuron.

**Fig 8.**
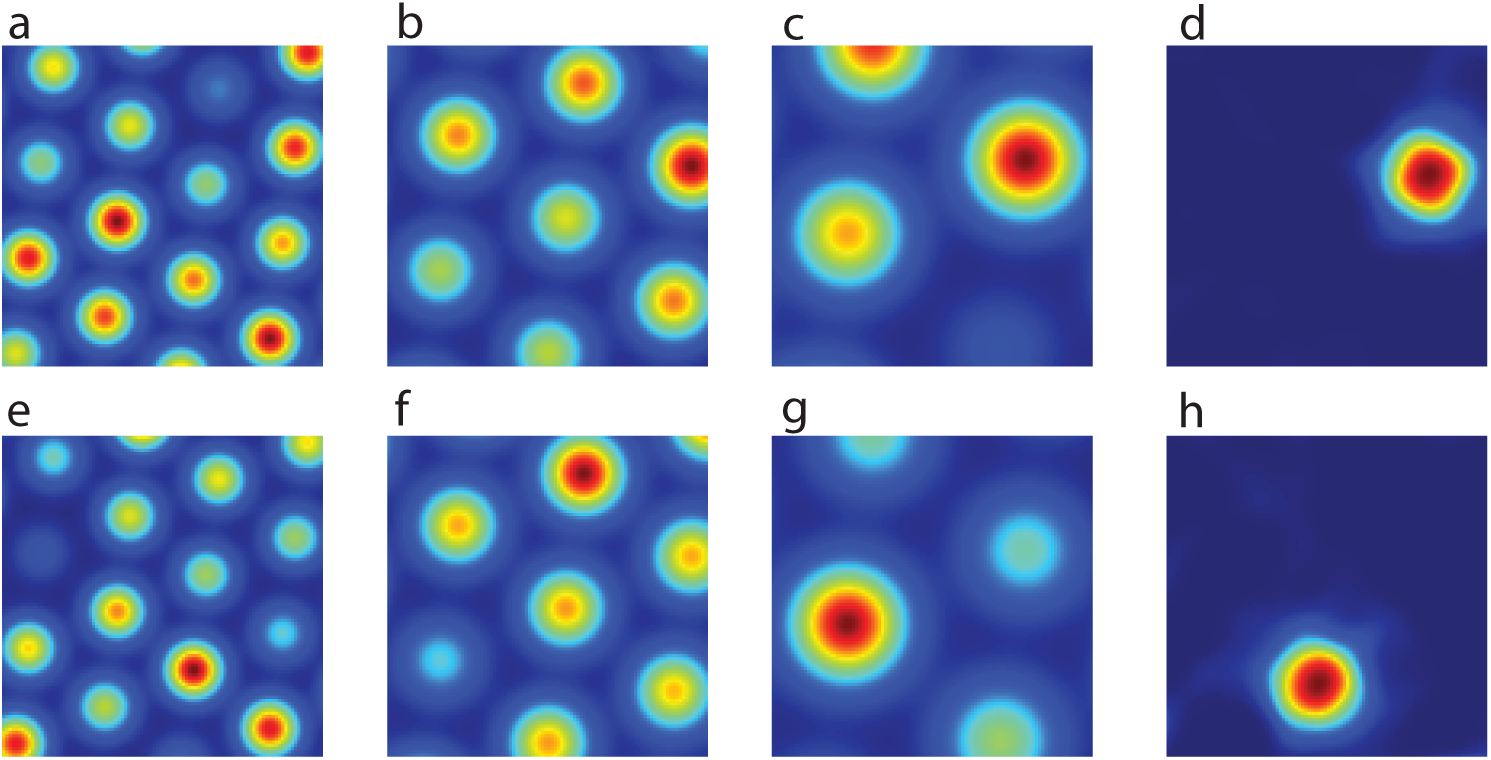
Rate-remapping of grid cells sufficient to globally remap a place cell. a-c: Example simulated grid cells from 3 modules with imposed field-to-field variability. **d**: A place cell with only input from grid cells of the type in **a-c** (25 from each of the 3 modules) and a constant external input. **e-g**: Same cells as in **b** but with different field amplitudes. This manipulation has been accomplished chemogenetically in rats [28]. Here the peak rates of the fields were sampled from a Gaussian distribution with a mean of 1 and standard deviation of 0.3. **h**: The same place cell as in **d** but with the input grid cells rate-remapped (as shown in **e-g**). Thus with the same connections and constant external input a place cell can globally remap with just a rate-remapping of the grid cells. Here the grid cell selectivity was simulated with Gaussian bumps centered on the vertices of a hexagonal lattice while the firing rate of the place cell is determined by the input with a logistic activation function.

To consider the consequences of this variability in firing field amplitudes within the context of the continuous attractor model, we devised two recurrent network models of grid cells, explicitly including both excitatory and inhibitory populations, and using patterned (fully factorized) or random (partially factorized) connectivity from grid cells to interneurons, respectively. We considered these networks operating under three different conditions: only receiving input from place-like input, only receiving a velocity dependent signal, or receiving only a constant input which we used for studying the drift. While the components necessary for the velocity input are found locally in the MEC [2,4], the MEC has also been shown to receive extensive input from the hippocampus as well as other areas with discrete spatial selectivity such as the postrhinal cortex [40,41]. Indeed, even during hippocampal inactivation, in which grid cells loose their periodic selectivity, stable discrete spatial selectivity in the form of single fields as well as increased directional selectivity has been demonstrated in grid cells [42].

We found that the fully factorized model could generate stable grid cells by only integrating velocity input and employing only a very small population of interneurons, e.g. approximately 5 % of the size of the excitatory population. In this condition, the grid cells did not show noticeable firing field amplitude variability. By adding a place-like signal via random connectivity to the excitatory cells, we obtained grid cells that exhibited amplitude variability in their fields, similar to what we established to occur in the real data. The degree of this variability could be controlled by diluting the spatial input to the grid cells or adjusting the variance of the strength of the connections from the place-like input.

Regardless of the type of input, the interneurons in the fully factorized models demonstrated periodic spatial coding with high grid scores, albeit with different kinds of firing patterns such as the “anti-grid” pattern. Interestingly, the interneurons maintained high grid scores and relatively low field-to-field variability while grid cells became more variable with increasingly variable input connections. This may indicate that the input connection strengths to an interneuron are distributed quite uniformly among grid cells with similar spatial phases in this model, such that the field-to-field variations of single grid cells often cancel approximately when summing over several cells of the same phase.

With only velocity input, the partially factorized model failed to generate the periodic firing pattern associated with grid cells. When replacing velocity input with a place-like input, however, the excitatory neurons acquired grid cell firing with realistically varying firing field amplitudes, but with amplitude variance increasing significantly faster than in the fully factorized model as a function of the variance of the place-like input connection strengths. The interneurons in this model, lacking the patterned input connectivity required to maintain their firing patterns while grid field amplitudes fluctuate, were strongly impacted by the variance of the connections from the place selective cells, and lost their periodicity as the variance increased.

In order to understand why this network failed to generate grid cells with only velocity input, we evaluated the attractor drift in the models, and observed that while there was a rapid exponential decrease in drift with the number of interneurons in the fully factorized models, even with a moderate number of interneurons (figure 7), the decrease was much slower and did not reach the same levels even at an order of magnitude more interneurons in the partially factorized model.

We interpret these results by proposing two possible scenarios for the underlying network that generated grid cells. Since both models are able to generate grid cells firing patterns in excitatory neurons with the field amplitude variability found in experimental data, both can be used as models for the generation of grid cells. However, they are distinguished by their predictions for the firing patterns of interneurons and their response to different types of input, and may thus differ in their compatibility with experimental data as we discuss further below.

The selectivity of interneurons in the partially factorized model (figure 4, right) resembled the selectivity of the majority of recorded local interneurons in Buetfering et al. [27]. However, the model required place selective input in order to generate grid cells, a prediction consistent with developmental data showing that place cell selectivity appears before grid cells [43].

The fully factorized model (figure 4, left), managed to generate grid cells with only velocity input, and firing field amplitude variations could be observed in the network when place-like input was added. However, this model also generated periodic spatial selectivity in the interneurons. Although the majority of the interneurons recorded in Buetfering et al. [27] did not show periodic firing, a few of them still did, which is consistent with the fact that the fully factorized model can generate the hexagonal pattern of the grid cells by integrating velocity input using a very small number of interneurons compared to the total size of the network. An interesting experimental proposal is therefore to investigate how many of the interneurons in the MEC are actually involved in generating grid cells, e.g. by looking at the fraction of interneurons that project back to grid cells, and seeing if these are among the few interneurons that show periodic firing, or by optogenetically manipulating a fraction of interneurons and seeing how it affects the grid cells’ periodic firing.

It should be mentioned that the use of place-like input in generating grid cells is not an entirely new idea. The so-called adaptation model originally developed by Kropff and Treves [21], and related models [44,45], consider the entorhinal network not as a fixed code with discrete translation symmetry built for path integration, but one that learns and adapts to different environments. In these models, external spatially modulated input provides an important role in the formation and maintenance of the grid pattern. Here, we have not discussed mechanisms through which the place-like input connectivity to the grid network may be formed through development or synaptic plasticity while the animal navigates a new environment. However, mechanisms such as those proposed by Kropff and Treves [21] might admit the establishment of proper place-like connectivity to the grid cells. Furthermore, though it has yet to be studied, it seems plausible that incorporating the observed field amplitude variability within such a framework may also be achieved by assuming that the spatial input can vary on a time scale that is faster than the learning of the connectivity required for the pattern to form.

The model presented here shares the much of the benefits of the adaptation model, such as stability to the types quenched noise [36,46] that plague models based on integrating velocity while still uses attractor dynamics enabling, for example, firing fields to appear for cells where external spatial input was removed or not provided. It is still an open question, however, what mechanisms for path integration could exist that are consistent with emerging properties, such as variability among grid cell firing fields.

## 5 Materials and Methods

### 5.1 Data

The 373 identified cells used in this analysis were selected from recordings of rats 13855, 13960, 14147, 15314, and 15708 in Stensola et al. [30], and details about the data acquisition can be found in that reference. These rats had multisite (hyperdrive) tetrode implants, and the recordings were carried out in square open-field environments of either 100 × 100 cm (rat 13855) or 150 × 150 cm (others).

### 5.2 Position filtering

The position time series, which were recorded at sampling rate 25 Hz, were smoothed to reduce jitter using a low-pass Hann window FIR filter with design cutoff frequency 2.0 Hz and kernel support of 13 taps, or approximately 0.5 s.

For each sample, the filter kernel was renormalized to compensate for missing samples within the kernel duration. In practice, this is accomplished by applying the filter to the original sequence but with missing samples replaced by zeros, and then to an indicator sequence with a value of 1 at every valid sample and 0 at every missing sample. The elementwise ratio of these two sequences is the final filtered sequence. This approach automatically interpolates across gaps shorter than the kernel duration, and is the simplest example of normalized convolution [47].

All experiments provided two sequences of position samples, corresponding to LEDs located in slightly different positions on the animal. The animal position was taken to be the mean of the two (filtered) sequences.

The speed at each sample was computed as the average speed over the surrounding 13 samples, by dividing the trajectory length with the elapsed time within this window. This was used to apply a speed filter: only samples where the speed exceeded 5 cm/s were kept.

### 5.3 Rate maps

Spike locations *ξ_j_* = (*ξ_j_*, *χ_j_*) at spike times *τ_j_* were found by linearly interpolating the position time series (*t_i_*,*x_i_*,*y_i_*) at the spike times. The spatial firing rate at location *x* = (*x*,*y*) was defined as the ratio of kernel density estimates of the spatial spike frequency and the occupancy,

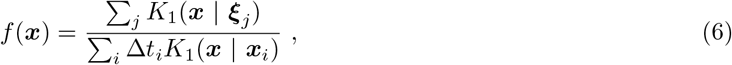

where the sums are taken over the spikes and position samples, respectively. Here, Δ*t_i_* = (*t*_*i*+1_ − *t*_*i*−1_)/2 ≈ 1/*F_s_* where *F_s_* = 25 Hz is the target sampling frequency A triweight kernel was used,

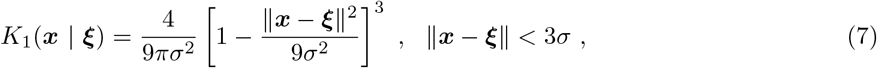

with bandwidth *σ* = 4.2 cm for 150 cm environments and *σ* = 2.8 cm for 100 cm environments.

The triweight kernel has compact support, so we avoid the infinite extrapolation entailed by a Gaussian. It is also a smooth, bell-shaped bump, that eliminates some problems that in our experience tend to show up when using finite kernels that go non-smoothly to zero (such as the uniform or Epanechnikov kernels), in particular that the ratios of such KDEs tend to blow up along the edges of the support.

Model simulations returned a continuous firing rate rather than discrete spikes, providing a combined position-rate time series (*t_i_*, *x_i_*, *y_i_*, *r_i_*). Rate maps were created by considering each rate sample as a fractional spike and otherwise proceeding as with spike trains,

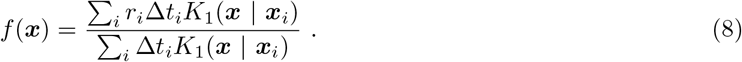

All rate maps were discretized for further analysis by sampling f(x) on a uniform square grid with bin centers *x_ij_*. We write *f_ij_* = *f*(*x_ij_*) Unless otherwise stated, a 5050-bin grid was used.

### 5.4 Correlograms

Spatial rate correlograms were computed from discretized rate maps using a Pearson product-moment correlation coefficient of overlapping regions of the rate maps at all integer-bin spatial lags. The cross correlogram of *ρ* rate maps *f* and *g* can be expressed as:

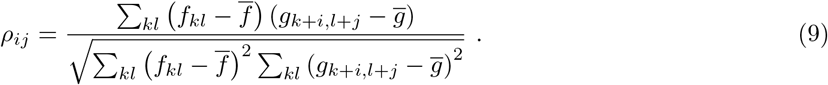

All sums were taken over the overlapping region of the rate maps at the given lag *i*, *j*. A single global mean 
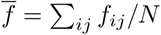
was used at all lags (here *N* is the number of bins in *f_ij_*, usually 50 × 50 = 2500).

### 5.5 Scores and selection

The grid score of a cell was computed from its spatial rate autocorrelogram by considering the angular Pearson autocorrelation of the autocorrelogram at lags of 30°, 60°, 90°, 120°, 150°, as well as ±3° and ±6° offsets from these angles. Only bins closer to the center than a radius *u* were included in the angular autocorrelations, and the grid score at a particular *u* was defined as the difference between the average of the maximum correlations around 60° and 120° lags and the average of the minimum correlations around the 30°, 90° and 150° lags:

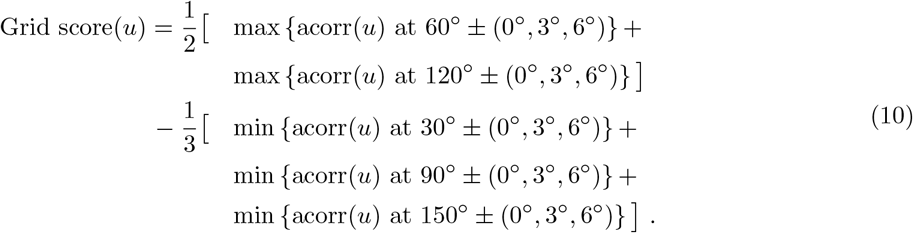

The final grid score of the cell was defined as the maximum of Grid score(*u*) over values of u ranging from four bin widths (i.e. *u* = 12 cm for 50 × 50 bins in a 150 cm box) to the full width of the experimental environment (half the width of the autocorrelogram), computed at intervals of one bin width.

We also calculated the spatial sparsity score defined by Buetfering et al. [27] (adapted from Skaggs et al. [48]) to quantify the degree of spatial tuning in the rate map:

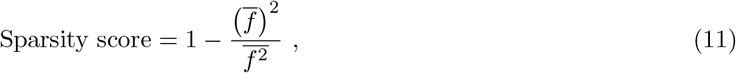

where *f* denotes the firing rate of the cell and the bars indicate an occupancy weighted spatial mean, i.e. a mean where each bin is weighted by the value of the occupancy map in that bin. The occupancy map is the denominator in equation 6.

The selection of cells for further analysis was done using a combination of these scores:

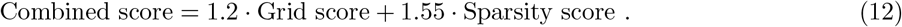

Only cells for which the combined score exceeded 1:0 were included in the further analysis. The purpose of this selection was to only include cells for which it was possible to make out clear and distinct firing fields in the spatial rate maps (quantified by the sparsity score) and for which these fields were laid out in a grid-like pattern (quantified by the grid score). The form of the combined score allows for an additive tradeoff between the two properties, while rejecting cells for which both scores are low. The exact weights were determined by looking for an appropriate cutoff line through grid score-sparsity space (see Supplementary figure 1) that separated cells that were subjectively deemed too noisy or disordered from the others.

In order to exclude suspected multiple recordings of the same cell, an additional selection filter was applied. For each pair of cells recorded from the same animal, one cell was excluded if the following criteria were satisfied:

1. The cells were recorded on the same tetrode.
2. The cells were either
  - recorded closer in depth than 75 *μ*m (for datasets where tetrode depth estimates for each session were available), or
  - recorded closer in time than nine days (for other datasets).
3. The Pearson correlation of the cell rate maps at zero spatial lag exceeded 0:8.

Note that the issue of possible repeats in these datasets was also addressed in the original reference [30, Supplementary Materials and Methods], and we have only used data that was kept at that time.

### 5.6 Lattice vectors

The spatial rate autocorrelograms were separated into segments associated with different peaks using a watershed algorithm. The algorithm is seeded with all the local maxima in the correlograms, and works by interpreting the negative of the correlogram as a topographical map and virtually “flooding” the image from imaginary water sources at the seeds. Thus each source is assigned a basin, separated from neighboring basins along the watershed lines where water from different sources meet as the water level is increased. Segmentation was only applied to regions of the correlogram exceeding a correlation threshold *ρ_min_*, where we used *ρ_min_* = 0.1 as a default value and made adjustments for certain cells in order to properly separate the central peak and the inner ring of six peaks around it. We always discarded all segments associated with a seed in an edge bin of the correlogram, but only after performing the segmentation with such seeds included.

The exact location of a peak was defined as the center of mass of the associated segment of the correlogram.

The inner six peaks provide three vectors that would ideally correspond to primitive lattice vectors of the grid pattern of the cell. (Due to the inversion symmetry of the autocorrelogram, there are only three unique vectors up to a trivial sign difference.) However, while a lattice can be defined by any two of the vectors extracted from the autocorrelogram, the third vector will in general not conform exactly to the same lattice. Therefore, we derived a projection from the peak vectors to a corresponding set of consistent lattice vectors.

Let *p_i_*, *i* = 1, 2, 3 be three unique peak vectors from the autocorrelogram, numbered in successive order around the origin. In addition, let *a_i_* be a set of vectors that define a consistent lattice and are closely approximated by the *p_i_*. In order to define a lattice, the *a_i_* must satisfy

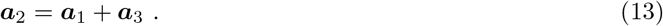

The vector *a_2_* must then be close to both *p_2_* weighted average of the two: *a_2_* = (*μp*_2_ + *p*_1_+ *p*_3_)/(*μ* + 1) Extending this reasoning to *a_1_* and *a_3_*, we arrive at the transformation

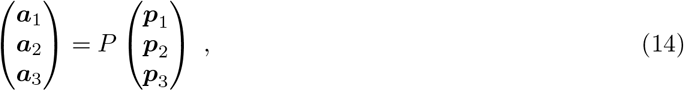

with

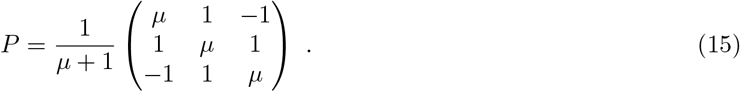

This transformation is by construction equivalent to the identity whenever the *p_i_* already satisfy 13. To guarantee that the *a_i_* always define a consistent lattice, it is therefore sufficient to require that P is a projection matrix, i.e. *P^2^* = *P*. This gives the unique solution *μ* = 2. The effect of this projection is shown in Supplementary figure 2.

In practice, we found it more convenient to work with all six peaks in the inner ring in the autocorrelogram. The projection derived above is straightforward to extend to that case, and the result is

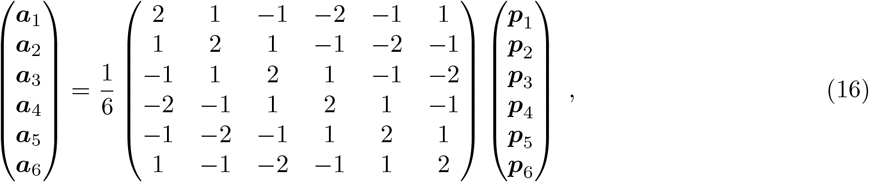

where the vectors are numbered in successive order around the origin.

In the following, we write the Cartesian components of *a_i_* as (*a_x,i_*, *a_y,i_*)

### 5.7 Deformation ellipse

To quantify the deformation of the grid cell firing pattern from a perfect triangular lattice, and also to define the spacing of the grid pattern, we fitted an ellipse through the endpoints of the lattice vectors *a_i_*. This fitting problem has an exact solution, which we derive here.

A general conic section can be written as an implicit quadratic equation in two variables:

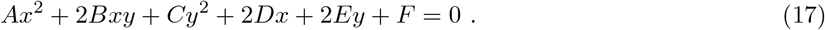

Scaling the six parameters *A, B, C, D, E, F* by a common factor will not change the conic, leaving five degrees of freedom for size, shape and orientation. The conic will be an ellipse if *B_2_* − *AC* < 0.

For *N* points (*x_i_*, *y_i_*), *i* = 1,…,*N*, we define the *N* × 6 design matrix *M* as

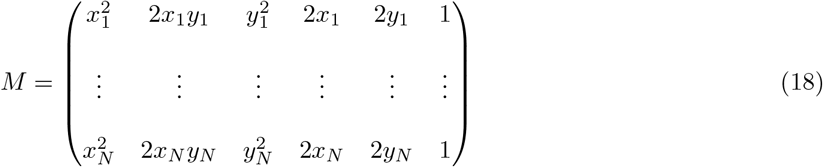

The conic section with parameter vector *θ* = (*A, B, C, D, E, F*)^T^ passes through all the points if *M θ* = 0. When *M* has rank 5, and hence a one-dimensional null space, this equation uniquely determines θ up to a scaling factor, and hence there is a unique conic section passing through all *N* points. In this case the system can be solved via singular value decomposition of *M* - the right-singular vector corresponding to the vanishing singular value is the parameter vector *θ*.

If we construct *M* from the lattice vectors *a_i_* = (*a_x,i_*, *a_y,i_*), and use the fact that *a*_*i*+3_ = −*a_i_*, *i* = 1, 2, 3 (assuming cyclically wrapping indices), we can write

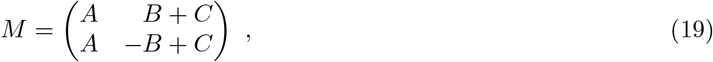

which is row equivalent to

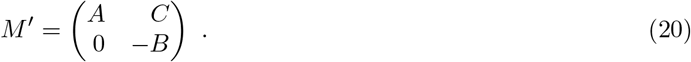

Here,

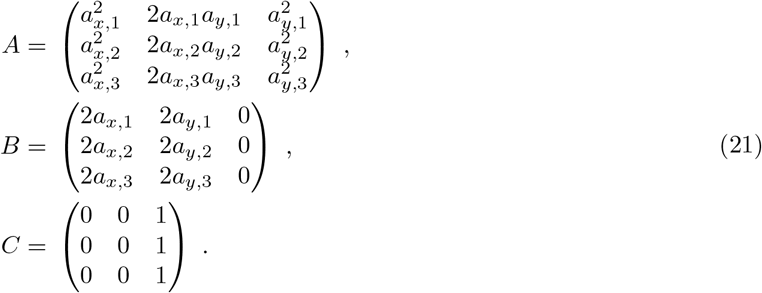

Provided that *a_1_*, *a_2_* and *a_3_* are not all parallel, we see that rank *B* = 2. As long as none of them are parallel, we also find that *A* is full-rank (the easiest way to see this is to divide row *i* in *A* by 
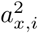
, and consider the linear independence of the resulting rows). Hence, rank [*A C*] = 3, and thus rank *M* = 5. This is exactly the condition required to solve exactly for the conic section passing through the lattice vector endpoints. Note that this derivation only relies on inversion symmetry among the a_i_, so the conic can just as easily be fitted through the unprojected peak vectors *p_i_*^1^.

We have not shown that the fitted conic will be an ellipse, but the inversion symmetry among the *a_i_* rules out the possibility of obtaining a parabola, and a hyperbola will not be obtained from points that are vertices of a convex polygon. Both these properties should always be satisfied for the inner ring of peaks in a grid cell autocorrelograms.

From the analytic geometry parametrization *θ* = (*A, B, C, D, E, F*)^T^ one can derive the equivalent canonical parameters (*x_c_*, *y_c_*, *a*, *b*, *θ*) of the ellipse (see e.g. [49]). Here, (*x_c_*, *y_c_*) are the coordinates of the center of the ellipse, *a* and *b* are the lengths of the semi-major and semi-minor axes, and θ ∊ [−π/2, π/2] is the angle between the major axis of the ellipse and the coordinate *x*-axis. Defining the translated and rotated coordinates

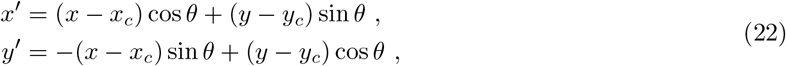

the ellipse satisfies

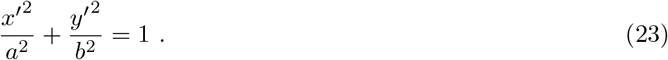

### 5.8 Grid spacing

We defined the grid spacing λ of a cell as the geometric mean of the lengths of the semiaxes of the deformation ellipse, 
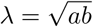
 Thus defined, λ is the radius of a circle with the same area as the ellipse, and it is therefore preserved under any area-preserving deformation of the grid pattern, such as the shearing described in Stensola et al. [50].

An ellipse fitted through a ring of grid cell autocorrelogram peaks, and a circle with the same area, are shown in Supplementary figure 3.

### 5.9 Grid cell feature space

We defined a space of grid cell features in which Euclidean distance was used as a proxy for the similarity of the cells’ firing patterns. The feature vector of a cell can be written as

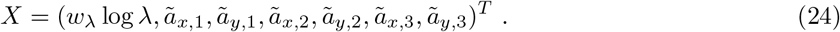

Here, we define 
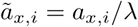
and similarly for 
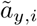
 These are dimensionless versions of the grid lattice vectors, and characterize the orientation and deformation of the grid pattern independently of the spacing. This way, the grid pattern shape and size are decoupled in feature space, and their relative importance is tuned by the weight *w_λ_*. The grid spacing λ only enters through its logarithm, so that all differences and distances in feature space are dimensionless and independent of the measurement units used (log λ_2_ − log λ_1_ = log λ_2_/λ_1_).

### 5.10 Module clustering

Modules were identified by detecting clusters in the grid cell feature space using the mean-shift clustering algorithm [51]. The algorithm works by randomly choosing a candidate centroid, and iteratively updating it until convergence by replacing it with the centroid of all points within distance *h*. This is repeated for a number of different starting locations, and after eliminating near-duplicates, each resulting centroid corresponds to a cluster in the dataset. A sample which was only visited by (i.e. within distance *h* of) centroid trajectories that converged to the same final centroid is taken to be part of the corresponding cluster, while remaining points are declared outliers. The algorithm takes a single parameter h, the bandwidth, and determines the number of clusters automatically.

We discarded outliers and clusters with less than five cells, and accepted the remaining clusters as grid cell modules. Using the feature space weight *w_λ_* = 3.3 and the mean shift bandwidth to *h* = 0.35, we reproduced the modules from Stensola et al. [30] in all animals^2^.

In order to assert the stability of the clusters, the clustering was repeated 10 times on each dataset. The clustering was identical in all 10 runs on all datasets when using the mentioned parameters.

### 5.11 Average firing field shape

We used a small region around the central peak in the autocorrelogram as as a model for the shape and size of a typical firing field of the cell. We defined this region using a separate correlation threshold *ρ*_field_, and modeled the firing field shape as a 2D Gaussian bump with variance Σ^2^ = −*A*/2π log(*ρ*_field_) where *A* is the area of the region around the central peak where autocorrelation exceeds *ρ_ff_*. This definition is designed to match the value of the Gaussian at distance 
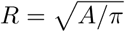
 from the origin, relative to the peak value, to the correlation threshold *ρ*_field_, i.e. 

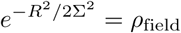

We used the threshold *ρ*_field_ = 0.55 for all cells. We defined the radius of the average firing field as *u _f_* = 1.6Σ.

### 5.12 Idealized spatial tuning profile

We derived an idealized spatial tuning profile from a grid cell by tessellating space with the lattice defined by the primitive lattice vectors *a_i_*, and placing a circular Gaussian bump with variance Σ^2^ at each node. The lattice was translated to the location that maximized the zero-lag cross correlation between the rate map of the actual cell and the idealized profile, and scaled to have the same mean firing rate as the recorded cell. The tuning profile is meant to represent an idealization of the spatial tuning of the real grid cell, with the same lattice parameters, spatial phase, firing field size and mean firing rate, but with firing fields that are identical.

### 5.13 Firing field detection

Firing field centers were detected in grid cell rate maps using the same watershed segmentation algorithm used to detect peaks in the autocorrelograms (see section 5.6). The detection was performed on rate maps created with 1.5 times the bandwidth of the ordinary rate maps, i.e. *σ* = 6.3 cm for 150 cm environments and *σ* = 4.2 cm for 100 cm environments. Segmentation was applied only to regions of these rate maps where the firing rate exceeded the 25th percentile of the distribution of rates in the rate map.

Candidate firing fields were defined as circular regions of radius *u_f_* around the detected firing field centers. The amplitude *r* of a field was defined as the number of spikes recorded within this field region, divided by the time spent there.

A final filtering step to distinguish true firing fields from possible noise blobs was performed by comparing the candidate fields to the firing fields in the idealized spatial tuning profile derived from the rate map, using the same radius *u_f_* for the idealized and candidate fields. We computed the overlap fraction q of each pair of a candidate and an idealized field, defined as the ratio of the overlapping area of the fields to the field area π*u*^2^_*f*_. Pairs with *q* > 0.25 were sorted by the product *rq*, where *r* is the amplitude of the candidate field, and unique pairs were kept starting from the top of this list. Thus false positives among detected fields were eliminated, and each detected field was associated with an idealized counterpart.

For rate maps created from sampled spikes, firing fields were detected in the exact same way, using the same idealized fields as for the corresponding real rate map (i.e. no new idealized spatial tuning profile was created for the synthetic rate map). Corresponding fields in the real and synthetic rate maps were identified by association with the same idealized field, and only fields detected in both the real rate map and all synthetic rate maps from the same cell were used further analysis.

For rate maps created from model simulations, no lattice parameters were extracted from the autocorrelogram and no idealized tuning profile was created. Hence, candidate fields were used directly without any filtering. This is unproblematic, since model cell rate maps are not noisy.

Supplementary figure 5 shows the detected firing fields and corresponding amplitudes for a selection of real grid cells and an accompanying synthetic rate map per real cell.

### 5.14 Sampled spike trains

We used the idealized spatial tuning to sample spike trains corresponding to this spatial tuning and the real animal trajectory, assuming Poisson spiking statistics and no covariates except spatial location. To this end, the real trajectory of the rat was divided into Δ*t* = 5 ms bins, and the target firing rate *f_i_* at in bin *i*, centered on time *t_i_*, was found by evaluating the tuning profile at the corresponding location (*x_i_*, *y_i_*). In each bin, a Bernoulli trial was performed with success probability *f_i_*Δ*t*, and if successful a spike was placed at *t_i_*.

Rate maps were computed from these spike trains in the same way as for real cells.

### 5.15 Statistical analysis of firing field amplitudes

Only cells for which 3 or more fields were detected (counting only fields detected in the real rate map and all synthetic rate maps derived from it) were used in the statistical analysis of firing field amplitudes. We computed the sample coefficient of variation of the amplitudes of a cell as

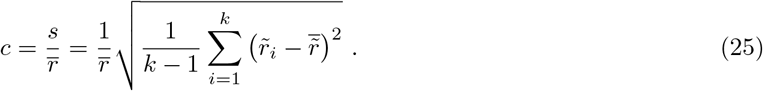

Here, k is the number of fields, *r_i_*, *i* = 1,…,*k* are the amplitudes of the fields, and 
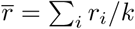
 is the mean amplitude. The usual sample variance is given by *s*^2^.

To compare the temporal stability of the field amplitudes with the field-to-field variations, we extracted firing field amplitudes using only data from the first and second halves of the experiment, respectively (using the same fields as for the full-length data). We denote these amplitudes as *r_ij_*, where the index *i* runs over the *k* firing fields and *j* runs over the partitions 1,2. First, we define the following sample means:

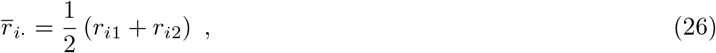

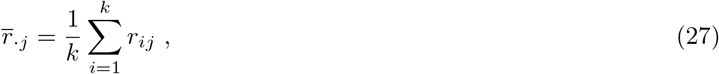

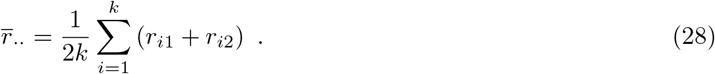

The between-field variability is defined as,

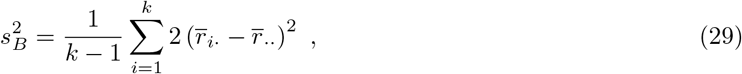

and the within-field variability as

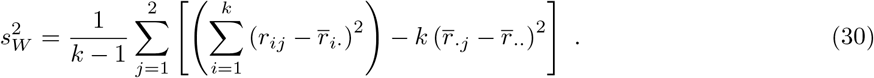

The second term in equation 30 is a repeated-measures correction included to remove contributions to the within-field variability due to a coherent shift in amplitudes between the two halves of the experiment. We do this because we are only interested in capturing random within-field fluctuations that may explain the measured between-field variability, not coherent variations that would not contribute to this and may even be due to experimental artifacts such as as the cell not being detected by the tetrode for parts of the experiment. We compensate by subtracting n ‒ 1 from the degrees of freedom of the estimator (i.e. dividing by (*k* − 1)(2 − 1) = *k* − 1 instead of *k*(2 − 1) = *k*), such that *s*^2^_*W*_ remains unbiased.

A key point here is that if all the r_ij_ are independent and identically distributed, both *s*^2^_*B*_ and *s*^2^_*W*_ are unbiased estimators of the variance of this distribution. Hence, if *s*^2^_*B*_ is significantly larger than *s*^2^_*W*_, this may indicate that there is a component of variability between the field amplitudes that cannot be explained by fluctuations that affect between-and within-field variabilities alike. This is made quantitative via the

-statistic,

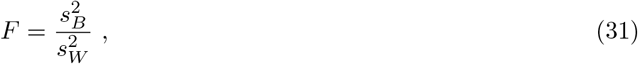

which may be familiar from one-way ANOVA. For each cell, we computed a *p*-value based on this statistic by ranking it against the same statistic computed from the 1000 simulated spike trains associated with the cell. (Notably, we did not use critical values of the *F*-distribution to get the *p*-values, as this would be entail unjustified normality assumptions)

We also define dimensionless coefficients of variability, useful for plotting and comparison:

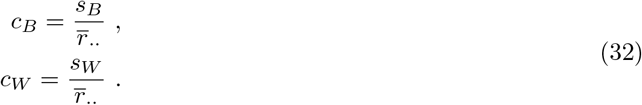

The *F*-statistic can be expressed in terms of these quantities:

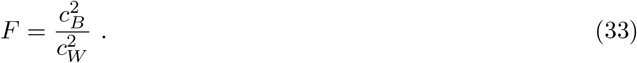

### 5.16 Two population continuous attractor network model

To investigate the firing properties of interneurons in the two population model with place-like external input, we constructed networks consisting of 32 × 28 grid cells and 200 interneurons for the fully factorized model, and 4480 interneurons for the partially factorized model. The activity was translated during exploration via place-like input from 180 × 180 units each encoding one unique position of the environment and projecting to a single grid cell. The values of the non-zero connections from the place-like neurons was drawn from a Gaussian distribution with mean 0.5, truncated at 0.0. The values for the standard deviation ranged from *σ* = 0 to *σ* = 5 for the fully factorized model, and *σ* = 0 to *σ* = 1 for the partially factorized model. The path of the animal in the simulation was taken from two adjacent 10 minute recordings and resampled into 10 ms bins.

In the models to check the drift of the network, we used 16 × 13, 32 × 28 and 64 × 55 grid cells in the fully factorized case and 32 × 28 for the partially factorized model. The number of interneurons was chosen as a fraction of the total grid cell population. In each of these, the neurons were assumed to lie on a twisted torus [29] with connectivity as described in the main text and *R*_min_ = 0.5 × *N*_y_ and *R*_max_ = *N_y_*. The other parameters were chosen as *τ* = 10 ms, constant = 0.1, *W*_0_ = 0.1 and *l* = 0.

For the path integration model we went with a four bump model with 64 × 56 grid cells, 200 interneurons, *R*_min_ = 15, *R*_max_ = 20, *τ* = 10 ms, *l* = 4, *α* = 0.0075, **constant** = 0.1 and *W*_0_ = −0.05. As mentioned above, no place-like input were used for error correction during path integration. A 10 minute recording of the animal was resampled into 10ms bins and used to insure that the statistics of the movement were consistent with those of an actual animal.

### 5.17 NMF

The matrices **J** and **K** were first set to small random values and the updated iteratively according to the multiplicative learning rules

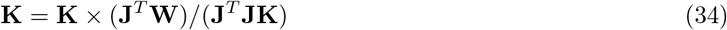

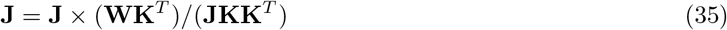

where the signs × and / represent element-wise multiplication and division, respectively. The above equations are iterated until convergence, i.e. the change in the value of the cost function 
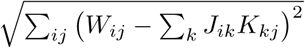
is below 0.0001.

## Acknowledgments

We thank Hanne and Tor Stensola as well as Edvard and May-Britt Moser for making the data available. We are also grateful to Edvard Moser for his comments on this work, and also Trygve Solstad for discussions at the early stages of this project. This work was supported by the Kavli Foundation, Norwegian Research Council (Centre of Excellence Scheme), and a Starr Foundation fellowship at the Simons Center for Systems Biology, IAS to YR.

## Supplementary Figures

**Supplementary fig 1.**
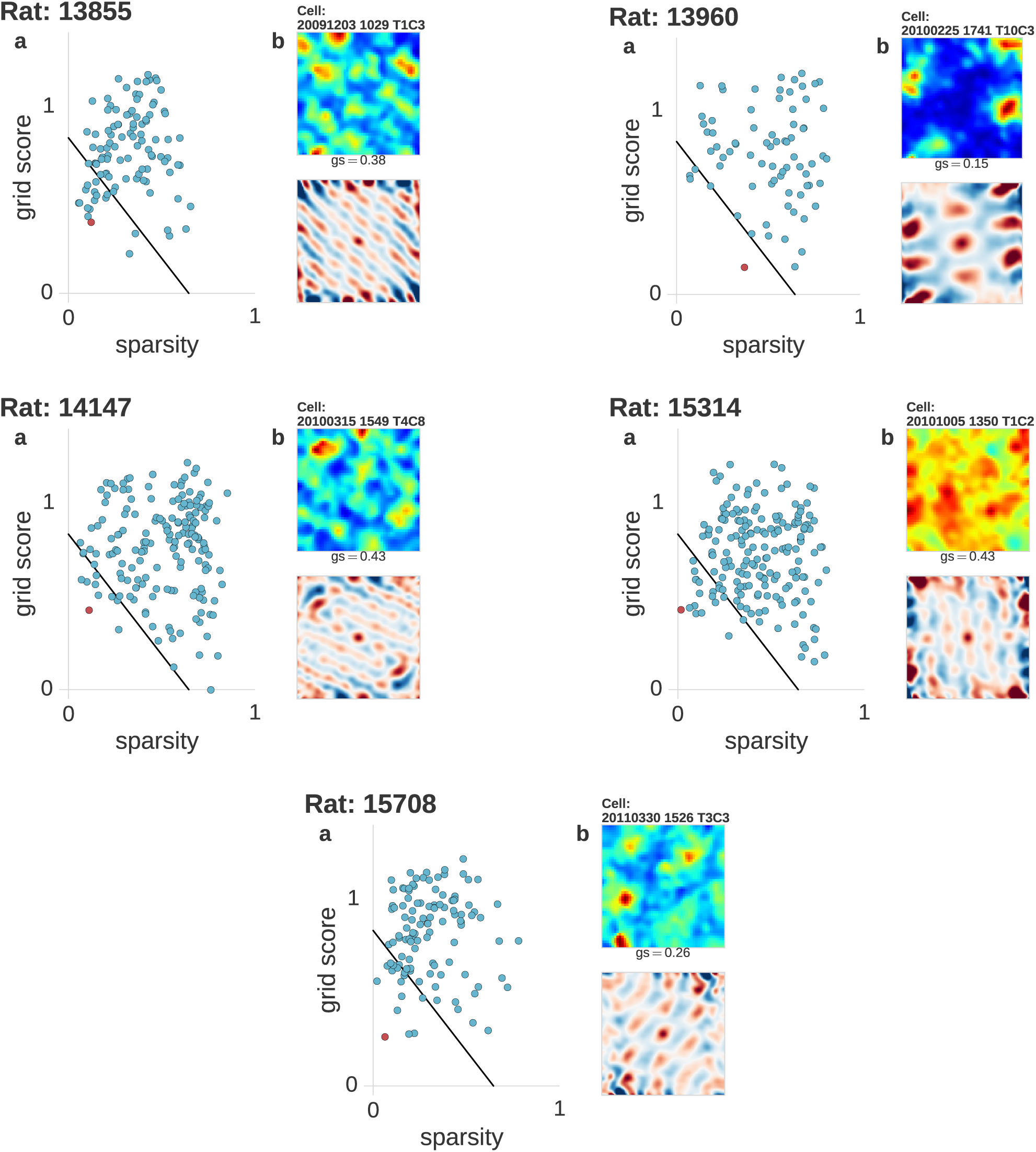
**Grid score-sparsity filtering. a**: All available cells scattered in sparsity-grid score space. The combined score cutoff defined in the Materials and Methods section is drawn with a black line: only cells above/to the right of the line were kept. The discarded cells tend to have noisy ratemaps, where firing fields and a clear triangular lattice may be difficult to make out. **b**: Ratemap and autocorrelogram of the cell with lowest combined score. This cell is also marked with a red dot in **a**. The grid score (gs) and sparsity (sp) of the cell are quoted below the ratemap.

**Supplementary fig 2.**
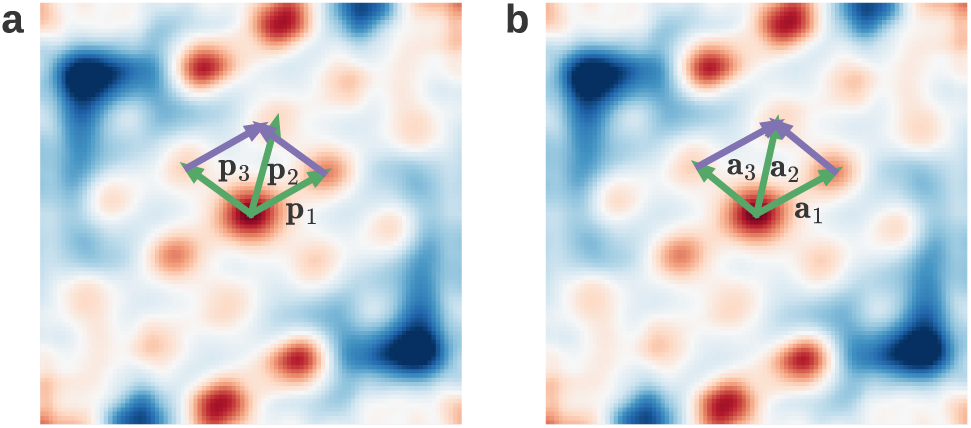
**Projection from peaks to lattice vectors. a**: A typical grid cell autocorrelogram with three detected peakcenters drawn as vectors *p*_1_, *p*_2_, *p*_3_ from the rigin. The purple vectors demonstrate that *p*_1_ + *p*_3_ ≠ *p*_2_. **b**: The same autocorrelogram, but now with vectors *a*_1_, *a*_2_, *a*_3_ that are related to *p*_1_, *p*_2_, *p*_3_ by the lattice projection derived in the Materials and Methods section. These vectors satisfy *a*_1_ + *a*_3_ = *a*_2_, and hence they define a consistent lattice.

**Supplementary fig 3.**
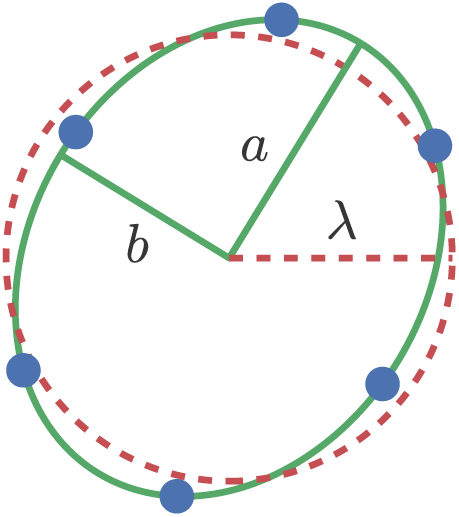
**Deformation ellipse and spacing of a grid pattern** The ellipse (green curve) has been fitted through a ring of autocorrelogram lattice points (blue dots) taken from the autocorrelogram. The semiaxes are labeled *a* and *b*. A circle with the same area as the ellipse is plotted concentrically (red dashed curve). The radius of this circle, λ = √*ab*, defines the grid spacing.

**Supplementary fig 4.**
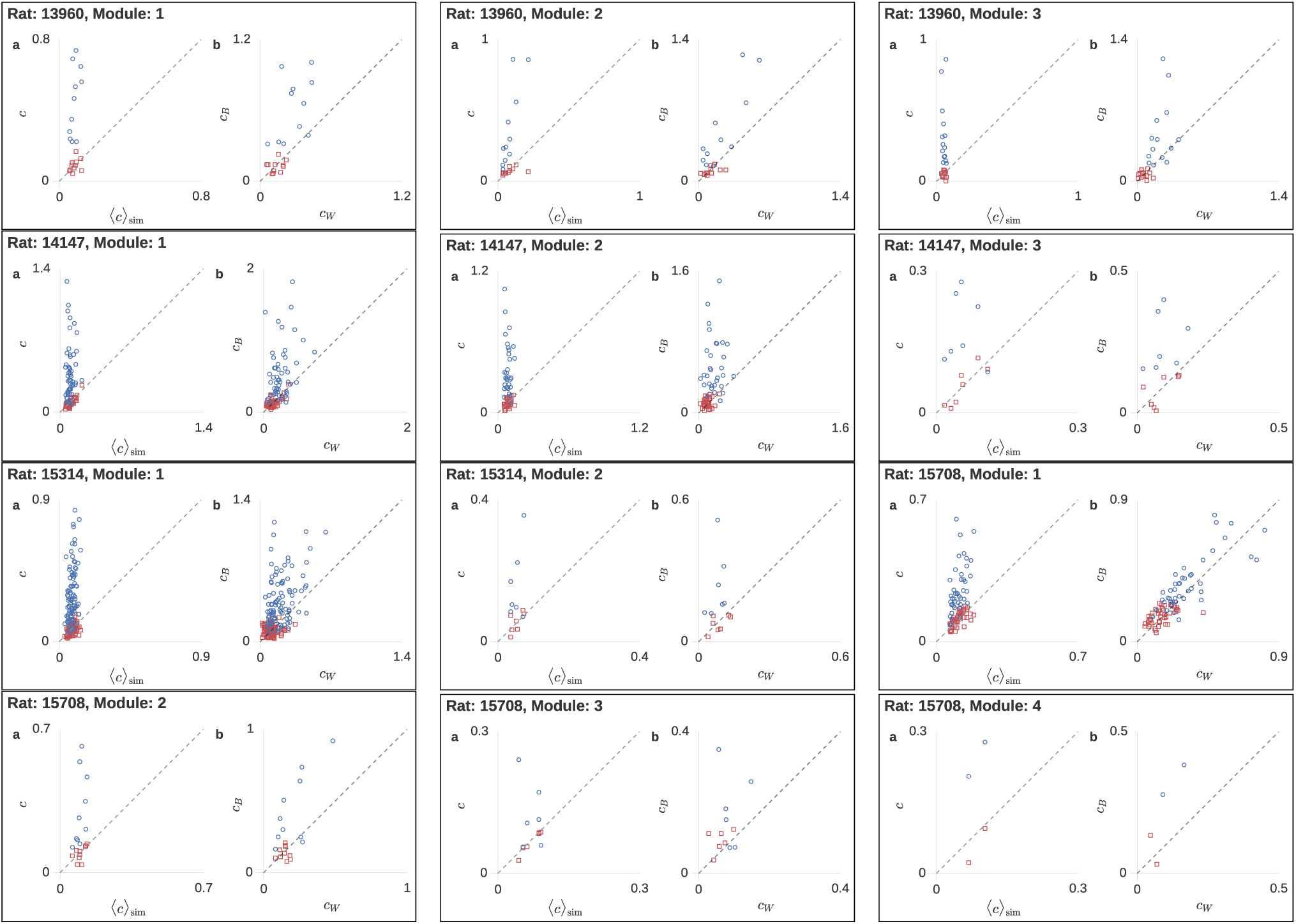
**Firing field amplitude variability. a**: Blue circles show the field amplitude coefficient of variation of each cell in a module, scattered against the mean coefficient of variation from the 1000 sampled spike trains corresponding to that cell. Red squares show the same quantity for one particular sampled spike train per real cell. **b**: Between-field coefficient of variability scattered against within-field coefficient of variability from the same real cells and synthetic spikes as in **a**. Variabilities are computed from field amplitudes measured in the first and second halves of the experimental sessions separately.

**Supplementary fig 5.**
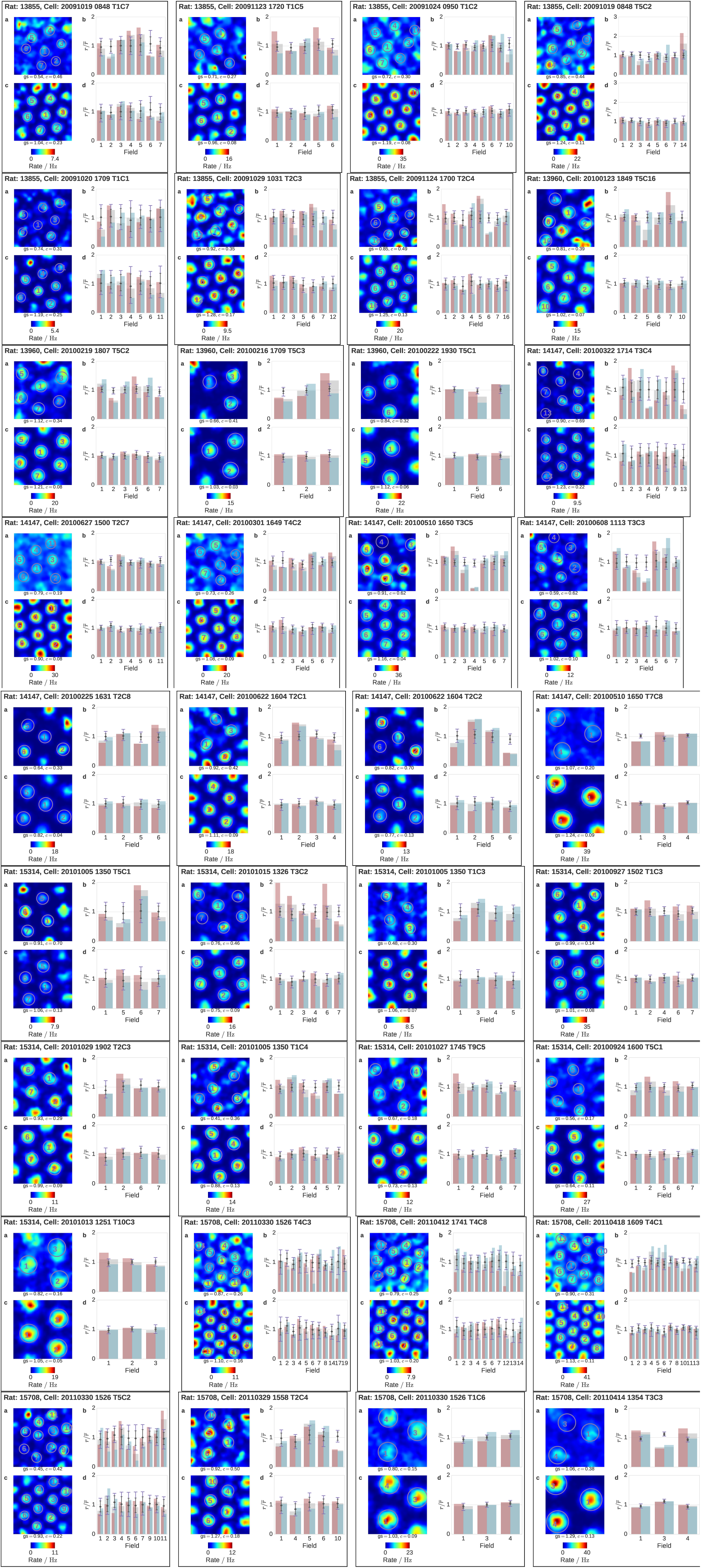
**Example grid cell ratemaps and firing field amplitudes, with simulated counterparts.** The cells used in this figure are a subset of the cells constituting the respective modules (approx. every 13th cell in each module). **a**: Grid cell rate map with detected firing fields circled and numbered. The grid score (gs), sparsity (sp) and field amplitude coefficient of variation (*c*) are quoted below the rate map. **b**: The normalized amplitude of each field in a is shown (gray wide bars in the background) together with the amplitudes measured in only the first and second halves of the session (red and light blue bars in the foreground). On top of this, markers and error bars visualize the distribution of normalized amplitudes from 1000 synthetic spike trains, with the black diamond showing the mean amplitude, the small black whiskers showing the central 95 % of the full-length amplitudes (to be compared with the gray wide bars), and the large purple whiskers showing the central 95 % of the half-length amplitudes (to be compared with the red and light blue narrow bars). For each cell and synthetic spike train, amplitudes are normalized to the mean of the full-length amplitudes from that cell or spike train when creating this plot. **c-d**: Equivalent to **a-b**, but using sampled spikes generated from the real animal trajectory and a tuning profile consisting of identical firing fields. Another 1000 synthetic spike trains like this were used to generate the error bars in **b** and **c**, and to perform the statistical analysis of the variability.

**Supplementary fig 6.**
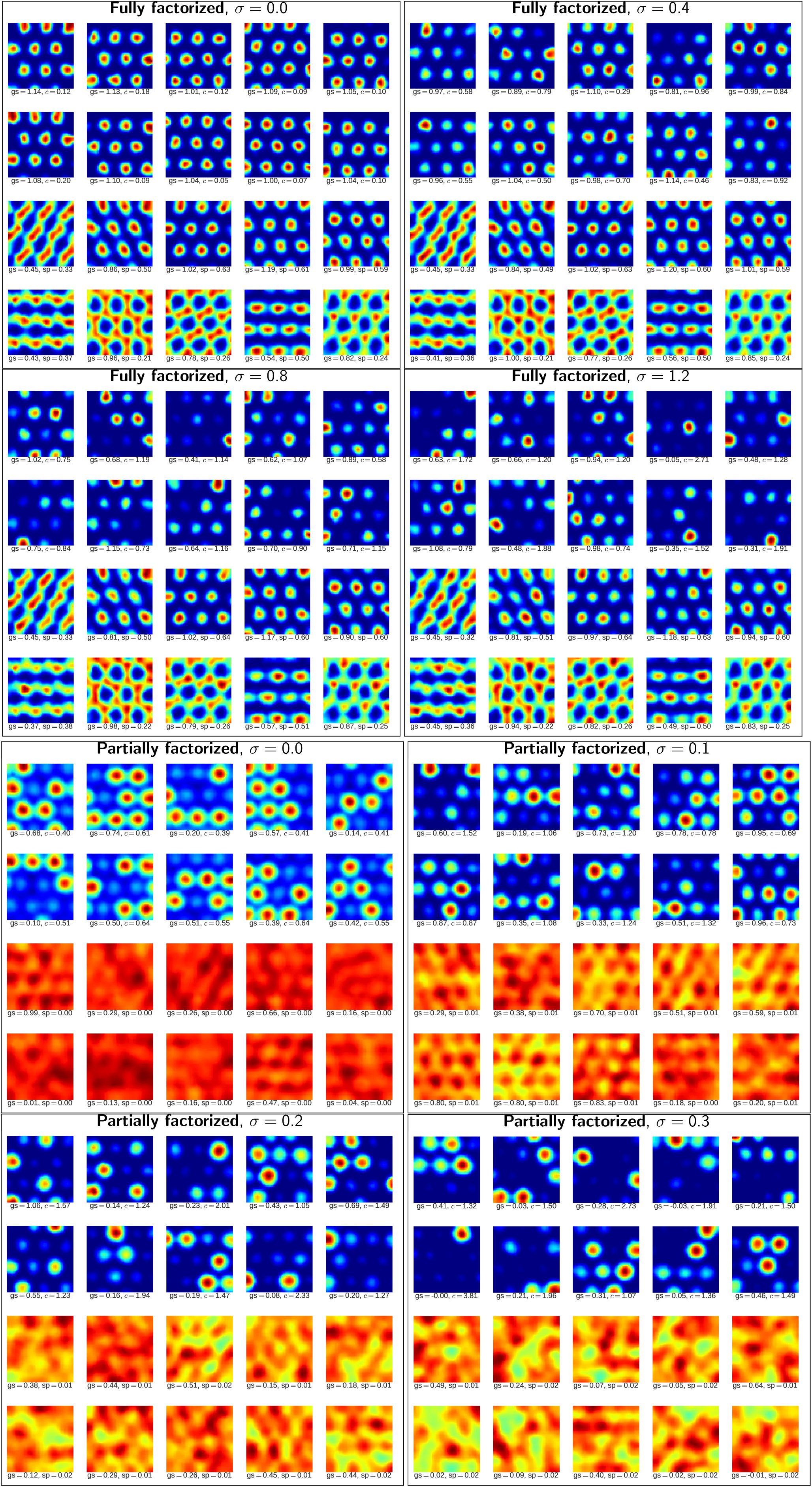
Additional examples of rate maps from the model simulation runs used in figure 4. In each frame, the two top rows are excitatory neurons (grid cells) and the two bottom rows are inhibitory neurons (interneurons).

**Supplementary fig S7.**
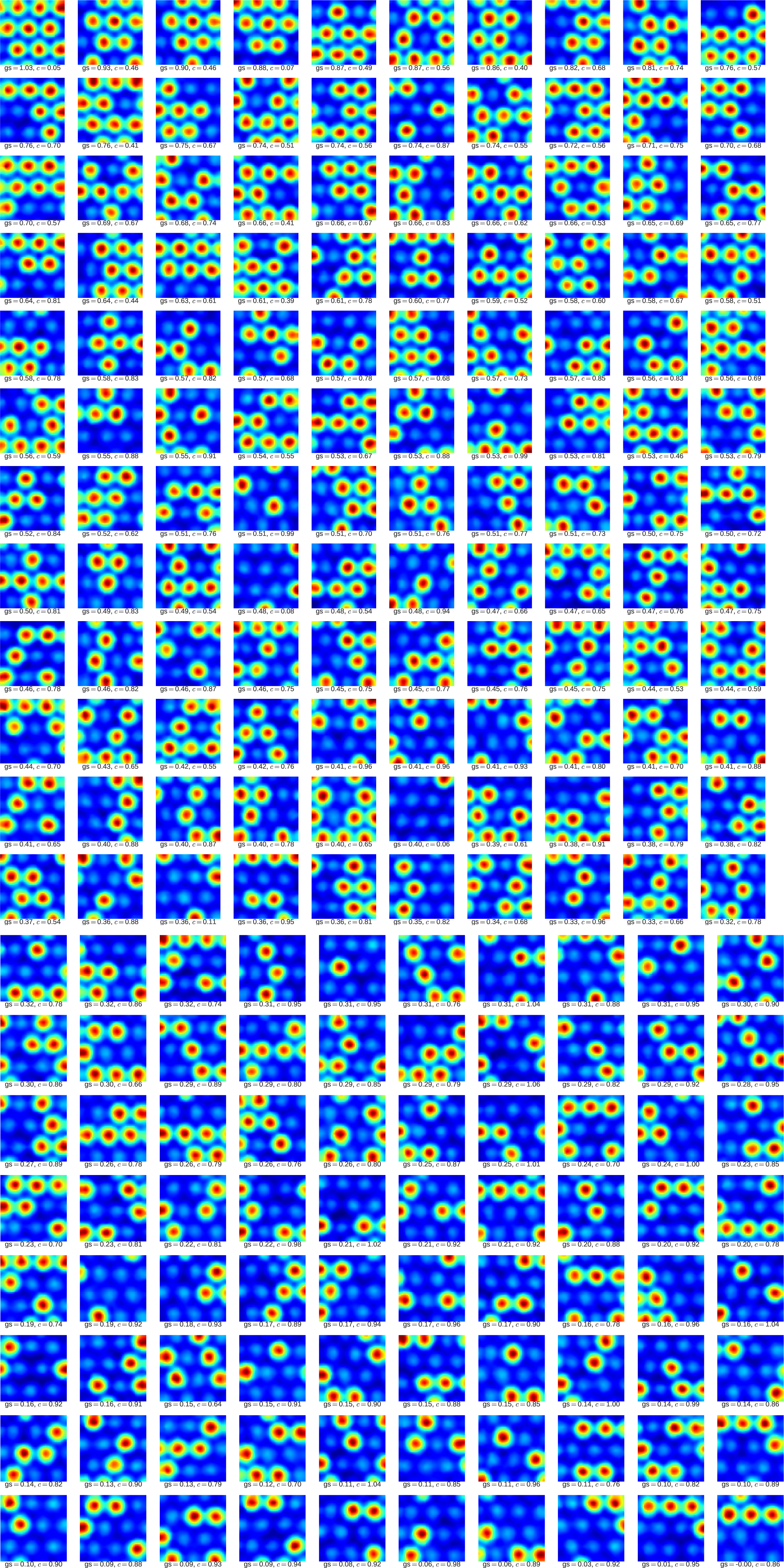
Ratemaps of all 200 excitatory neurons (grid cells) in a run of the partially factorized model with zero connectivity variance ( *σ* = 0) and dilution fraction *f_d_* = 0.5. Grid score (gs) and field amplitude coefficient of variation (*c*) quoted beneath each ratemap. The ratemaps are sorted by grid score, in decreasing order.

## Footnotes

1 What we have shown here is really that six points with inversion symmetry through a common center, or equivalently, a center and three peripheral points, uniquely define a conic section, as long as the points lie on three different rays through the center.

2 By this we mean that we found the same number of modules, with approximately the same average grid parameters, in all animals. Since we have filtered out a number of cells in previous steps, and our clustering algorithm generates outliers, the allocation of cells to modules was not identical.

